# Population-scale Characterization of the Oral Microbiome and Associations with Metabolic Health

**DOI:** 10.1101/2025.10.28.685004

**Authors:** Haochen Xue, Anastasia Godneva, Feilong Tang, Huifa Li, Yulong Li, Ming Hu, Ruobing Li, Jionglong Su, Eran Segal, Imran Razzak

## Abstract

The oral microbiome may capture system-specific information about host metabolic health, yet large-scale, multi-system evidence remains scarce. We analyzed 9,431 participants in the Human Phenotype Project (HPP), integrating buccal-swab oral whole-metagenome profiles with 44 metabolic measures spanning liver ultrasound, continuous glucose monitoring (CGM), and dual-energy X-ray absorptiometry (DXA). Using a microbiome-wide association study (MWAS) framework, we constructed a multilayer map across strains, gene families and pathways, revealing widespread associations: 213 strains, 124,603 gene families and 299 pathways were significantly associated with metabolic measures. Prioritizing the strongest and cross-phenotype signals, we identified multiple oral features with most significant associations to metabolic health. For example, acyl carrier protein (ACP) was associated with lower liver inflammation and reduced adiposity, whereas polyamine biosynthesis and ceramide ***α***-oxidation tracked higher glucose variability and adverse liver and adiposity phenotypes. Leveraging these MWAS-derived signals, we trained disease classification models using phenotype-selected oral features, which outperformed full-feature oral models across six metabolic diseases. These association signals were also robust in oral-health sensitivity analyses in HPP, and key BMI and waist-circumference associations directionally replicated at the genus level in an independent cohort (n = 20,293). Together, these findings provide a population-scale oral–metabolic association map and highlight the potential of oral microbial markers as non-invasive tools for metabolic risk stratification.

## 1 Introduction

The oral microbiome is a complex and dynamic ecosystem at the interface between the host and the environment, playing a vital role in maintaining both oral and systemic health [1–3]. Beyond its well-established involvement in caries and periodontitis [4–6], oral dysbiosis has been linked to cardiovascular, autoimmune, and metabolic disorders [7, 8]. Mounting evidence links alterations in the oral microbiome to metabolic conditions such as non-alcoholic fatty liver disease (NAFLD), type 2 diabetes, and obesity [9, 10]. Clinical and experimental studies further support the links between the oral microbiome and metabolic diseases. In liver disease, reduced oral microbial diversity and enrichment of genera such as Streptococcus, Porphyromonas, and Actinomyces have been reported [11], while phylum-level compositional shifts characterize NAFLD [12]. Similarly, in glucose metabolism disorder, increased abundance of opportunistic taxa has been observed in individuals with diabetes [13]. Most of these findings are predominantly compositional and descriptive, leaving the functional characterization of oral–metabolic associations largely unexplored.

Despite the substantial progress in oral microbiome research, two major limitations have constrained advances (i) most of the existing studies rely on 16S rRNA gene profiling to assess microbial diversity and taxonomic composition in metabolically healthy versus diseased individuals. However, 16S rRNA sequencing [14] provides limited taxonomic resolution and does not directly capture microbial genes or functional pathways. Consequently, strain-resolved and function-level (gene family and pathway) links to metabolic phenotypes remain difficult to map. (ii) recent studies are statistically underpowered due to small sample sizes and the prevalent use of binary case–control designs focused on single diseases. This narrow framework overlooks the complex, multisystem nature of metabolic health, limiting the biological and clinical relevance of current findings.

In this study, we address the gaps mentioned above by using the Human Phenotype Project (HPP), a large-scale longitudinal deep-phenotyping cohort of 9,431 individuals [15]. Using metagenomic profiling (MetaPhlAn4 and HUMAnN3), we profiled standardized oral metagenomes from buccal swabs and analyzed microbial features across strains, gene families, and pathways. Furthermore, HPP used advanced techniques to collect a wide array of multi-organ metabolic health phenotypes, including liver status via liver ultrasound, dynamic glucose homeostasis via continuous glucose monitoring (CGM), and visceral adipose tissue via dual-energy X-ray absorptiometry (DXA). Building on these large-scale and high-dimensional data, we performed microbiome-wide association studies (MWAS) framework [16, 17] to systematically investigate associations between oral microbial features and host metabolic phenotypes. This study leverages the large-scale HPP cohort to build a population-scale, multi-system association map linking oral microbial strains, gene families and pathways to host metabolic phenotypes, providing resource for future mechanistic and clinical studies.

## 2 Results

### 2.1 Study design and research population

The Human Phenotype Project cohort includes 9,431 participants recruited for the collection of oral microbiome and metabolic phenotypes. We collected liver metabolic health data using liver ultrasound, continuous glucose levels via CGM, and DXA-based body composition data. Data modalities are summarised in Fig. 1a, and Fig. 1b illustrates per-participant coverage and the sampling window. The summary statistics for key metabolic phenotypes are shown in Table 1, which characterizes that HPP is a middle-aged cohort (mean age ∼56 years) with a near-balanced sex distribution. Expected sex differences are apparent (women higher total body fat %; men higher VAT/lean mass and slightly higher mean CGM glucose and GMI), and the wide spread of liver, CGM, and DXA measures supports downstream association analyses. The HPP cohort is introduced in Method 4.1, and the detailed collection methods of phenotypes are described in Method 4.3.

**Fig. 1:**
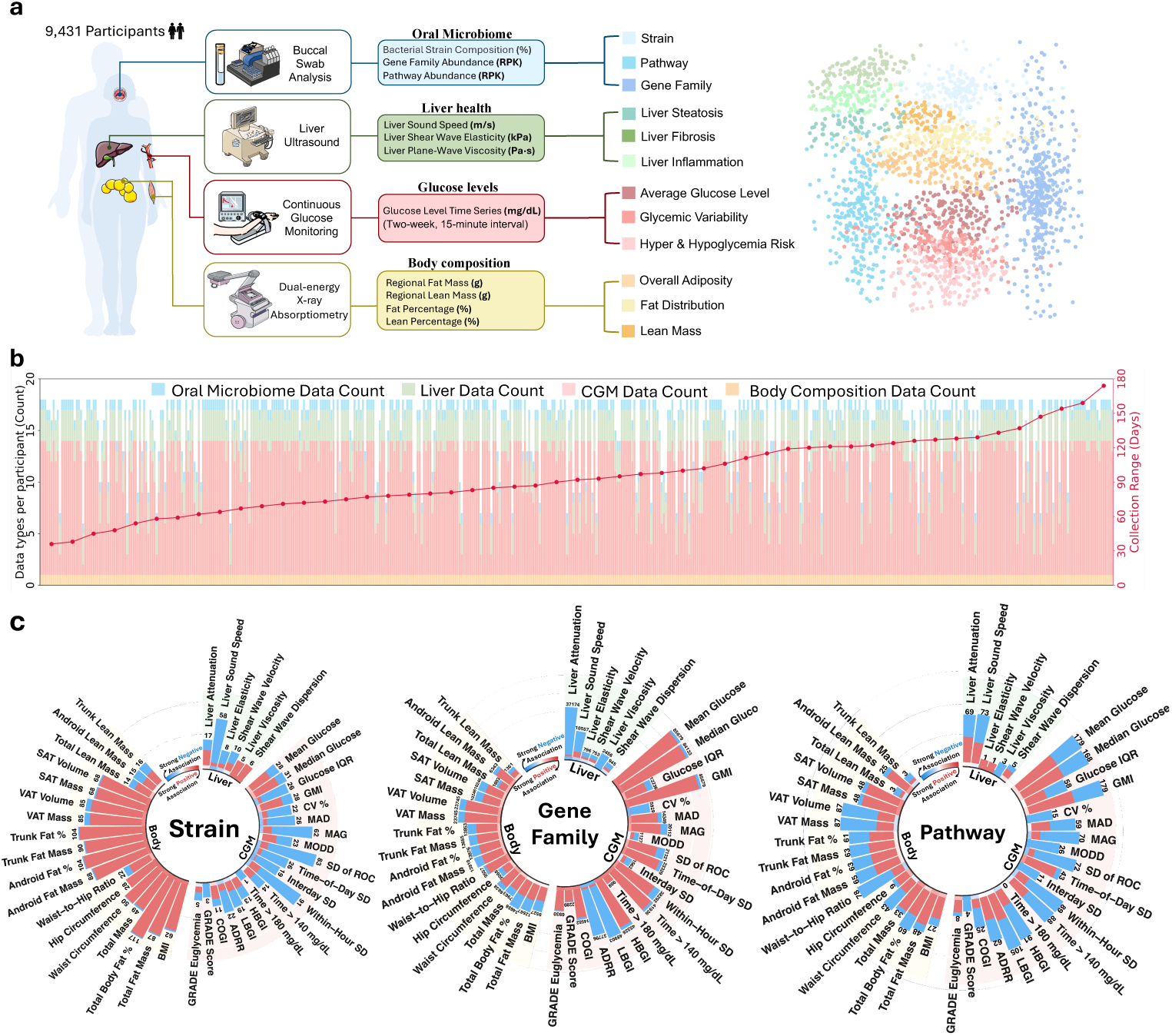
**a**, Data collected in the Human Phenotype Project (HPP; *n* = 9,431): buccal mucosal oral metagenomes (strains, gene family, and pathway), liver ultrasound (sound speed, elasticity, viscosity), continuous glucose monitoring (CGM; 15-min over 14 days), and dual-energy X-ray absorptiometry (DXA; body composition). Right: two-dimensional embedding of the 44 phenotypes, colored by physiological system. **b**, Coverage and timing for a random sample of 500 participants. Stacked bars show modality-specific phenotype counts; The red line (right y-axis) shows each participant’s data collection range, defined as the maximum absolute time difference between the oral-swab date and the liver ultrasound/CGM/DXA. **c**, Significant associations between oral microbial features and the 44 phenotypes at three levels (strain, gene family, pathway). Radial bars show counts per phenotype; colors indicate direction (red, positive; blue, negative). Significance: Bonferroni-corrected *P <* 0.05. The numeric counts underlying panel **c** are provided in Table S4.

**Table 1:** Summary statistics of key metabolic phenotypes. Values are mean (s.d.). Liver ultrasound phenotypes are grouped into liver steatosis, fibrosis and inflammation. CGM-derived phenotypes are grouped into average glucose level, glycemic variability, hyper-/hypoglycemia risk and overall glycemic control quality. Body composition phenotypes are grouped into overall adiposity, fat distribution and lean mass. The full list and extended statistics are provided in Supplementary Table S2.

| Metabolic Phenotype |  | Male<br>n=4474(47.4%) | Female<br>n=4957(52.6%) |
| --- | --- | --- | --- |
| **Age** |  | 55.49 (7.95) | 56.24 (8.01) |
| **Liver Health** |  |  |  |
| Liver Steatosis | Liver sound speed (m/s) | 1547.36 (23.07) | 1539.51 (23.69) |
|  | Liver attenuation (dB/cm/MHz) | 0.41 (0.10) | 0.41 (0.10) |
| Fibrosis | Liver Elasticity (kPa) | 5.43 (1.25) | 4.68 (0.94) |
|  | Shear Wave Velocity (m/s) | 1.34 (0.14) | 1.24 (0.12) |
| Inflammation | Liver Viscosity (Pa.s) | 1.70 (0.24) | 1.62 (0.27) |
|  | Shear Wave Dispersion ((m/s)/kHz) | 2.46 (0.45) | 2.32 (0.50) |
| **Glycemic Control (CGM)** |  |  |  |
| Average Glucose Level | Mean Glucose (mg/dL) | 97.35 (13.74) | 94.08 (11.23) |
|  | Glucose Management Indicator (GMI, %) | 5.64 (0.33) | 5.56 (0.27) |
| Glycemic Variability | Coefficient of Variation (CV, %) | 15.95 (4.18) | 16.15 (4.00) |
|  | Mean of Daily Differences (MODD, mg/dL) | 13.62 (4.93) | 12.88 (4.04) |
| Hyper/Hypoglycemia Risk | High Blood Glucose Index (HBGI) | 0.23 (0.94) | 0.14 (0.36) |
|  | Low Blood Glucose Index (LBGI) | 2.18 (1.91) | 2.57 (1.94) |
| Glycemic Control Quality | Composite Glycemic Index (COGI) | 0.89 (0.16) | 0.86 (0.17) |
|  | Diabetes Risk Assessment Equation (GRADE) | 1.37 (1.23) | 1.24 (0.85) |
| **Body Composition** |  |  |  |
| Overall Adiposity | Total Fat Mass (kg) | 23.16 (8.05) | 25.98 (8.56) |
|  | Total Body Fat (%) | 28.52 (6.32) | 38.71 (6.79) |
| Fat Distribution | Android Fat Mass (g) | 2327.65 (1105.68) | 2071.05 (963.79) |
|  | VAT Mass (g) | 1211.70 (748.66) | 647.77 (458.77) |
| Lean Mass | Total Lean Mass (kg) | 56.33 (6.82) | 39.67 (4.98) |
|  | Android Lean Mass (kg) | 4.07 (0.57) | 2.90 (0.43) |

To ensure data reliability, liver ultrasound was measured three times in a single day, and the average value was used for analysis. For CGM, interstitial fluid glucose levels were continuously monitored for two weeks, with readings taken every 15 minutes. For body composition data, 94.25% of participants had a single DXA measurement. For the 5.75% with multiple measurements (with time intervals ranging from 3 days to 22 months), data were processed in relation to the collection date of the oral microbiome sample. Specifically, if all of a participant’s DXA measurements fell within six months before or after their microbiome sample collection, the average of these measurements was used. Furthermore, for the correlation analysis, only metabolic phenotype data (liver ultrasound, CGM and DXA) collected within a six-month window centred on the oral-swab date were included.

Oral microbiome data were processed using MetaPhlAn 4 and HUMAnN 3.6 to characterize the taxonomic composition and functional potential. MetaPhlAn 4 provided the relative abundance of microbial taxa at all levels, from kingdom to strain. Subsequently, the abundance of gene families and pathways was quantified by HUMAnN 3.6 to reflect the functional profile of the microbial community. For subsequent correlation analysis, we used preprocessed strain-level abundances from MetaPhlAn 4 and functional data on gene families and pathways from HUMAnN 3.6, details in Methods 4.2.

Building on this time-aligned, cross-sectional dataset, we then (i) map oral microbial associations across 44 metabolic phenotypes (Results 2.2); (ii) prioritise cross-phenotype core markers (Results 2.3); (iii) perform proof-of-concept metabolic disease classification (Results 2.4); (iv) test robustness to oral-health confounding (Results 2.5); and (v) directionally replicate key oral–phenotype associations for BMI and waist circumference in an independent cohort (Results 2.6).

### 2.2 System-specific oral microbiome associations with metabolic phenotypes

Across 44 host phenotypes spanning liver, continuous glucose monitoring, and body composition, we analyzed 783 bacterial strains, 355,674 gene families, and 786 pathways. Among these, 213 strains (27.20%), 124,603 gene families (35.03%), and 299 pathways (38.04%) showed significant associations with at least one phenotype (Fig. 1c). Across all oral microbiome feature–phenotype pairs, positive associations predominated for strains (70.76%) and gene families (72.79%), whereas pathways exhibited a nearly balanced distribution (49.29% positive). Beyond these overall counts, two key patterns emerged. First, we observed extensive pleiotropy: most oral microbial features were linked to more than one host phenotype (90.20% of strains, 90.54% of gene families, and 94.98% of pathways). Second, we identified clear system-level specificity, with roughly half of the features (50.23% of strains, 59.29% of gene families, and 44.82% of pathways) showing associations restricted to a single metabolic domain—liver, CGM, or body composition. This coexistence of broad pleiotropy and system-level specificity formed the foundation for subsequent system-focused analyses.

Similarly, body composition exhibited the highest number of significant associations with strain at the system level, accounting for 63.68% of all significant strain–phenotype associations, compared to 30.57% for CGM-strain and 5.75% for the strain–liver associations. Furthermore, body also showed the highest association density, with a mean of 64.00 strains per phenotype, exceeding both liver (17.33) and CGM (27.74). Specifically, 46 strains were uniquely associated with body composition phenotypes and showed no significant associations with other systems. Among significant body–strain associations, positive relationships dominated (94.79%). Within body composition phenotypes, Total Body Fat% displayed the highest degree of association, with 117 strains significantly linked, of which 98.29% were positively associated. This strong positive was already evident before significance correction (87.4% positive) and remained pronounced after correction. At the genus level, 76.92% of the positively associated strains mapped to anaerobic, inflammation-associated genera (e.g., Prevotella, Actinomyces and Fusobacterium) that have been reported as enriched in obesity [18–20]. This positive dominant pattern was prevalent in all fat-related phenotypes: for measures of overall adiposity (e.g., BMI, Total Fat Mass), the mean proportion of positive associations was 96.62%; for indicators of fat distribution (e.g., Android Fat Mass, Trunk Fat Mass), the mean was 95.94%. A similar but attenuated trend was observed for the lean mass–related phenotypes, where the proportion of positive associations decreased to 60%. These results indicate that the strength of positive strain–host associations is strongly phenotype-dependent, being more pronounced for fat-related phenotypes than for lean mass.

At the functional level—encompassing gene families and pathways—the strongest associations shifted from body composition to CGM. Based on association counts, 70.20% of significant gene family associations and 57.41% of pathway associations were attributed to CGM. The CGM also exhibited the highest association density, with a mean of 29,951 significant associations per phenotype, exceeding liver and body composition (8,780 and 11,042, respectively). Moreover, the CGM also contained the largest number of system-specific signals, with 56,817 gene families and 107 pathways uniquely associated with CGM, markedly higher than the second-ranked body composition system (4,648 gene families and 16 pathways).

Given this strong system-specific enrichment, we next examined the directionality of CGM-associated features. Gene family associations were predominantly positive (78.66%), particularly for mean glucose and hyperglycemia metrics (95.34% positive), whereas hypoglycemia-weighted indices (LBGI, ADRR) were largely negative (4.08% positive). Within this overall positive framework, some divergence emerged among glycemic variability phenotypes: while most followed the positive trend, CV% exhibited far fewer associations and was overwhelmingly negative (1.97% positive). In comparison to gene families, CGM–pathway associations were more heterogeneous. For mean and median glucose, pathways mirrored gene-family directionality but with a lower positive fraction (61.38%). For variability phenotypes—including distributional (glucose IQR, CV%, MAD), short-term within-day (MAG, SD of ROC, within-hour SD), and between-day metrics (inter-day SD, MODD)—associations were predominantly negative, except for Time-of-Day SD, which remained largely positive (86.04%). This pattern aligns with the underlying metric definitions: Time-of-Day SD captures reproducible within-day glucose rhythms (e.g., pre-/post-prandial peaks, nocturnal troughs), whereas the other variability metrics primarily reflect stochastic short-term fluctuations or day-to-day instability. In contrast to CGM, the liver domain exhibited the fewest significant associations but showed clear pathology-specific patterns. For inflammation-related phenotypes, strain-level associations were uniformly positive, whereas pathway-level associations were predominantly negative (62.50%), with viscosity-related pathways being entirely negative. For fibrosis-related phenotypes, this directionality was reversed: strain-level associations were largely negative (66.67%), while both identified fibrosis-associated pathways (n = 2) showed positive associations. Together, these results indicate that, although the liver exhibited a lower overall association density, its microbial relationships were highly phenotype-specific and reflected distinct biological processes across inflammatory and fibrotic states.

To prioritize independent signals from the highly correlated oral gene families, we applied a two-step filtering strategy: a pre-association decorrelation filter applied before association testing, followed by post-association pruning based on the correlation structure (Methods 4.5). After pre-filtering and post-hoc clumping, the most robust and independent associations emerged for phenotypes reflecting overall glycemic status. For example, GMI retained 327 gene families after pre-filtering and 41 after clumping, while mean glucose retained 322 and 35, respectively—demonstrating substantial independence among signals. By contrast, certain phenotypes exhibited large unfiltered association counts but high redundancy. For instance, HBGI and LBGI were initially associated with 45,328 and 48,903 gene families, respectively; however, post-clumping reduced these to just 29 and 26 independent signals, revealing extensive collinearity within their underlying association networks.

### 2.3 Most significant associated oral strains, gene families, and pathways

Our analyses (Results 2.2) identified extensive and statistically significant associations between oral microbial features and host metabolic phenotypes, including 213 strains, 124,603 gene families, and 299 pathways. To identify the most informative microbial units, we applied a systematic selection strategy (Methods 4.6), identifying 15 strains, 15 gene families, and 13 pathways as core markers of host metabolic health. The associations for these core markers are summarized in Figs. 2, 3, and 4.

**Fig. 2:**
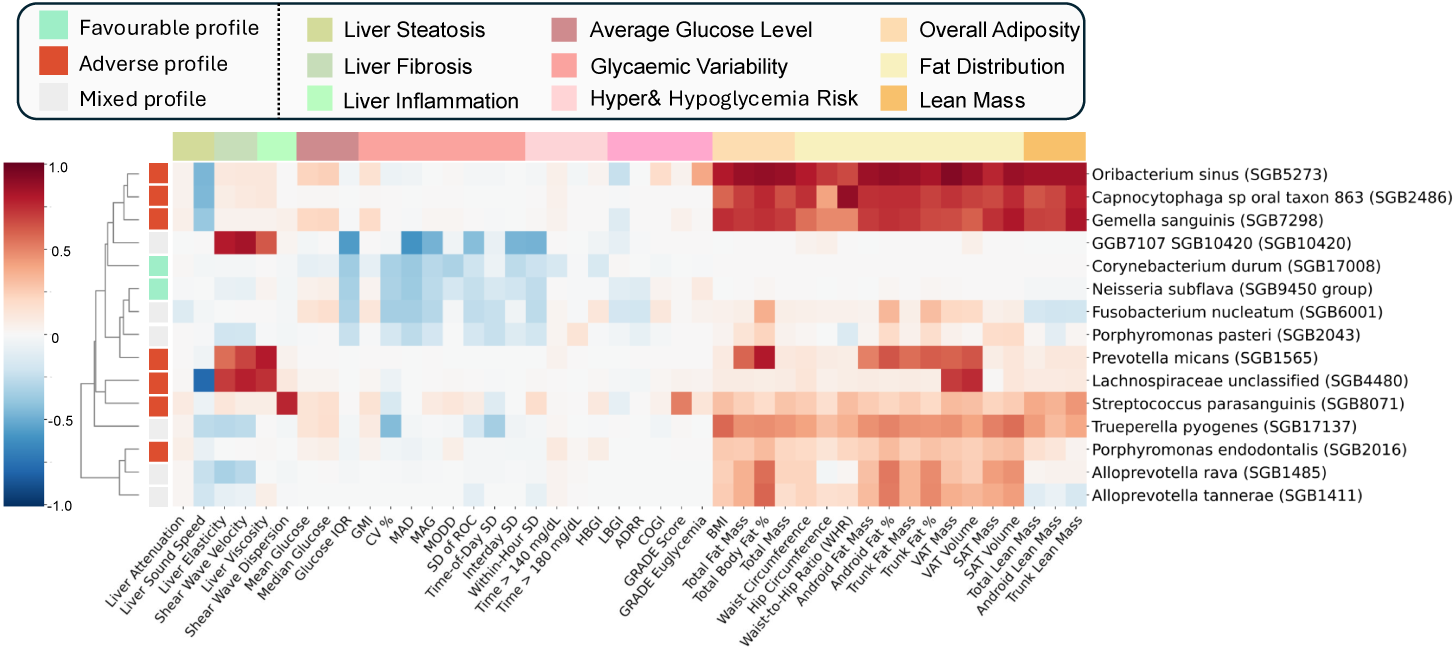
Heatmap illustrating the associations between 15 core oral microbial strains and 44 metabolic phenotypes. The selection method is detailed in Methods 4.6.

**Fig. 3:**
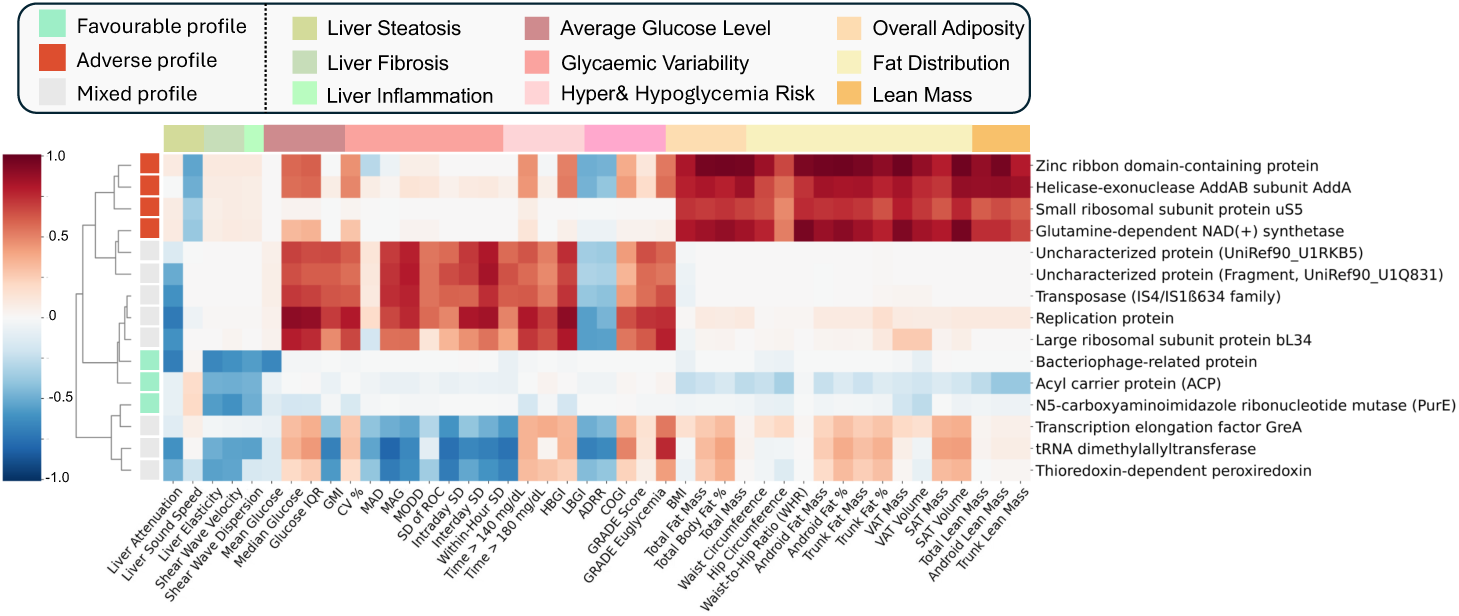
Heatmap illustrating the associations between 15 core oral microbial gene families and 44 metabolic phenotypes. The selection method is detailed in Methods 4.6.

Based on their cross-phenotype directions, we then grouped markers into three profiles: (1) favorable metabolic profile, indicating directions aligned with a healthier metabolic axis; (2) adverse metabolic profile, indicating directions aligned with a less healthy metabolic axis; and (3) mixed profile, indicating discordant directions across liver, CGM, and body composition phenotypes (Methods 4.7). Applying this same grouping to all Bonferroni-significant features yielded the following distributions: strains, 50/105/58; gene families, 17,782/44,648/62,173; pathways, 77/71/151 (favorable/adverse/mixed). The sections that follow examine the core strains, gene families, and pathways within each category in detail. Notably, a subset of core strains and pathways is consistent with a documented oral ecological gradient from oxygen-exposed mucosal/supragingival niches (e.g., Corynebacterium/Neisseria) to anaerobic, plaque-/subgingival-associated signatures (e.g., Porphyromonas/Prevotella/Oribacterium), a pattern that has been linked to gingival inflammation and increased biofilm burden in prior oral-biogeography and periodontal ecology studies [1, 4, 21]. Overall, our analyses provide a cohort-scale oral–metabolic association map resolved across strains, gene families, and pathways. Given the relative scarcity of strain-level data in the oral microbiome literature, we contextualize our findings using species- or genus-level reports where available and draw on non-oral studies for shared microbial genes and pathways when oral evidence is lacking.

#### 2.3.1 Significant strains

Among 15 selected most significant strains, 2 strains showed a favorable metabolic profile. Corynebacterium durum (SGB17008) associated with better glycemic control and lower CGM variability. Prior studies reported Corynebacterium durum (species-level) directionally concordant links to improved glycemic responses [10, 22]. Likewise, Neisseria subflava (SGB9450 cluster) was associated with more stable glucose and lower risk of hepatic inflammation and fibrosis, consistent with Neisseria subflava (species-level) identified as protective in metabolic dysfunction [23, 24].

In contrast, we also identified 7 strains with an adverse metabolic profile. Oribacterium sinus (SGB5273) and Prevotella micans (SGB1565) aligned with worse liver health and higher glucose and greater total/visceral adiposity; Lachnospiraceae taxon (SGB4480) showed a similar liver profile with increased adiposity/weight; Capnocytophaga taxon 863 (SGB2486) marked hepatic and body adiposity without CGM associations. In the CGM-anchored subset, Gemella sanguinis (SGB7298), Streptococcus parasanguinis (SGB8071) and Porphyromonas endodontalis (SGB2016) aligned with higher glucose and CGM variability, also increased body adiposity and liver steatosis. Where higher-rank reports exist, directions are consistent: Oribacterium sinus (species-level) has been linked to inflammatory responses under high-fat diet conditions [25, 26]; Prevotella micans (species-level) has been associated with greater fibrosis severity in NAFLD [27]; Lachnospiraceae taxa (family-level) have been associated with advanced fibrosis, increased liver stiffness and deteriorated glycemic profiles in gut cohorts [28]; Capnocytophaga (genus-level) has been implicated in NAFLD via oral–gut translocation [29]; Gemella sanguinis (species-level) has been reported as enriched in obesity cohorts [30, 31]; and Streptococcus parasanguinis (species-level) has been associated with hepatic steatosis, visceral-fat remodelling and impaired glucose regulation [32, 33]; Porphyromonas endodontalis (species-level) has been reported as enriched in periodontitis and obesity cohorts [34].

Finally, 6 strains exhibited a mixed profile. GGB7107 (SGB10420) showed lower CGM variability alongside higher liver inflammation and fibrosis. Fusobacterium nucleatum (SGB6001) followed a similar pattern, with lower CGM variability but higher adiposity and lower lean mass; this is consistent with Fusobacterium nucleatum (species-level) being linked to systemic metabolic disruption [35]. Porphyromonas pasteri (SGB2043) was associated with lower liver inflammation/fibrosis and lower CGM variability, but higher body fat and lower lean mass; this aligns with Porphyromonas pasteri (species-level) reports of increased liver inflammation/steatosis and higher body fat despite effects on glycemic regulation [36, 37]. Trueperella pyogenes (SGB17137) showed a similar split: more favorable liver indices and lower CGM variability, but higher glucose and higher body fat. Alloprevotella rava (SGB1485) and Alloprevotella tannerae (SGB1411) were associated with lower liver inflammation and fibrosis, but higher body / visceral adiposity and lower lean mass. These directions are consistent with Alloprevotella rava (species-level) and Alloprevotella tannerae (species-level) reports on inflammatory modulation alongside adiposity/liver changes [38–40]. To our knowledge, GGB7107 (SGB10420) and Trueperella pyogenes (SGB17137) had no associated reports related to metabolic health.

#### 2.3.2 Significant gene families

We identified gene families positively associated with metabolic health, primarily in core biosynthetic functions. For example, Acyl carrier protein (ACP; D4FQK5) associated with lower hepatic fat/inflammation and reduced visceral adiposity and body weight. Mechanistically, ACP serves as a core scaffold protein in the fatty acid biosynthesis pathway, and enhanced fatty acid metabolism can alleviate liver steatosis and inflammation [41]. In addition, we also observed favorable profile associations for uncharacterised genes, Bacteriophage-related protein (E7RXK3) from Lautropia mirabilis and N5-carboxyaminoimidazole ribonucleotide mutase (E6IZN5) from Streptococcus spp were both associated with improved liver health and glycemic indicators, to our knowledge, neither gene nor its source taxon has been linked to metabolic regulation.

A subset of gene families showed consistent adverse associations with metabolic health, mainly involved in translation, microbial proliferation, stress responses, and central metabolic regulation. Two ribosomal protein genes, the helicase-exonuclease AddAB subunit AddA (E4LF01) and the small ribosomal subunit protein uS5 (E6KU65), were associated with poorer liver function, increased adiposity, and increased body weight. The liver-related associations are supported by previous studies showing that ribosomal proteins are upregulated during dysbiosis and inflammation [42], and that increased ribosome biogenesis can directly promote hepatocyte proliferation and hepatic lipid accumulation [43, 44]. However, their associations with increased total adiposity and body weight are novel, with no prior evidence reported. Additional signals implicated regulators of metabolite levels and gene expression. Zinc-ribbon domain protein (D9RUZ0) showed adverse associations across liver, glycemic and adiposity traits; although direct evidence is lacking, its source genus Prevotella has been implicated in metabolic dysfunction [45–47]. Similarly, glutamine-dependent NAD^+^ synthetase (E6KUD8) was associated with overall worsening, consistent with data that supra-physiologic NAD^+^ promotes gluconeogenesis, impairs insulin signaling and enhances lipogenesis [48, 49].

We also observed a mixed profile: signals that were favorable for liver phenotypes but adverse for glycemic control. The unknown proteins U1RKB5 and U1Q831, IS4/IS1634 transposase (U1PZV2), a replication-associated protein (UPI0006850C96), and the large subunit ribosomal protein bL34 (U2QT17) were associated with lower liver fat, but higher glucose and greater glycemic variability. Genus-level context supports part of this pattern: U1RKB5 derives from Actinobaculum, which has been associated with improved liver and glycaemic profiles in NAFLD [50], and U1PZV2 from Actinomyces, enriched in MAFLD, encodes IS4/IS1634 transposases that may mobilize metabolic or drug-modifying genes, a potential route to glycemic effects [23, 51]; bL34 has also been implicated in metabolic syndrome and glucose dysregulation [52]. By contrast, U1Q831 and UPI0006850C96 currently lack functional or taxonomic links to host metabolism and merit follow-up. A second subset— Transcription elongation factor GreA (Z4WWF6), peroxiredoxin (J4VRI0) and tRNA-modification enzyme (J6H4J2), which combined lower liver fat/inflammation with higher glucose but lower variability, consistent with the impact of tRNA-modification pathways on glucose control [53, 54].

#### 2.3.3 Significant pathways

**Fig. 4:**
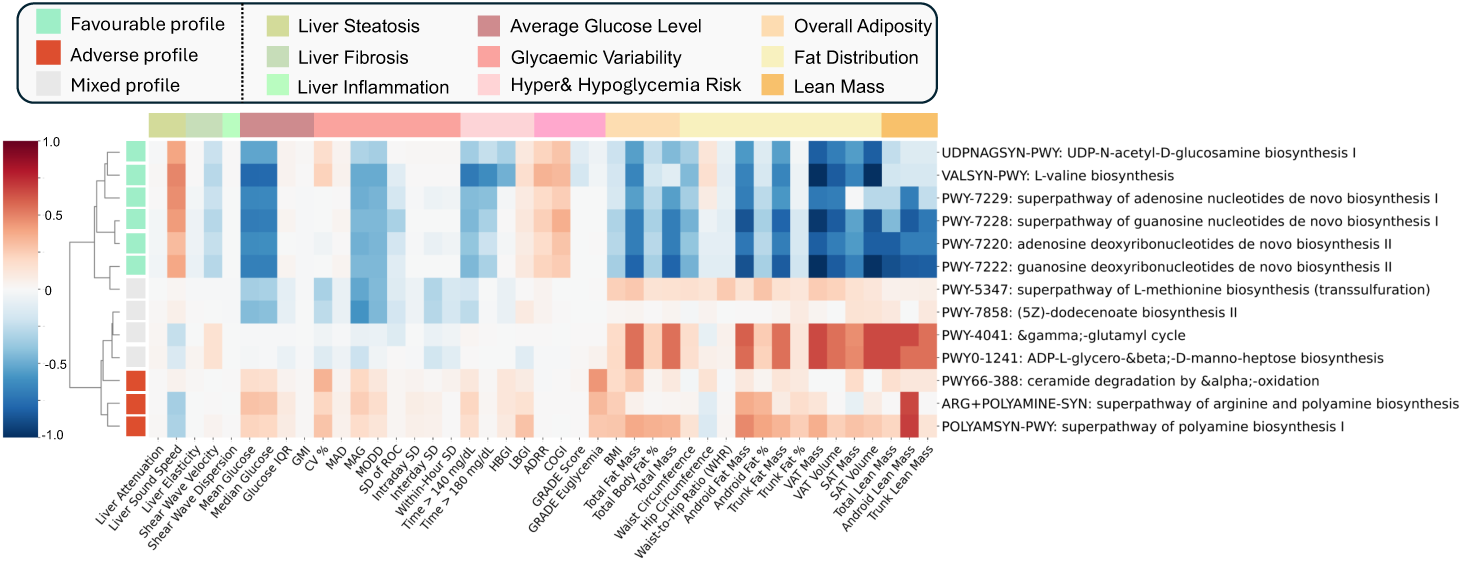
Heatmap illustrating the associations between 13 core oral microbial pathways and 44 metabolic phenotypes. The selection method is detailed in Methods 4.6.

We highlight 6 pathways associated with favorable metabolic profiles, particularly those involved in biosynthesis and energy metabolism. For example, UDP-N-acetyl-D-glucosamine biosynthesis (UDPNAGSYN-PWY) associated with better liver indices, more stable CGM and lower adiposity; non-oral evidence is directionally consistent for UDP-GlcNAc/hexosamine signalling [55]. L-valine biosynthesis (VALSYN-PWY) showed similar associations, aligning with BCAA supplementation benefits for glycemic control and hepatic lipid burden [56]. Additionally, Nucleotide de novo synthesis (tied to ATP/GTP) was broadly beneficial: purine routes (PWY-7229/-7220/-7222/-7228) correlated with better liver health, improved glycemic control and lower adiposity. Non-oral studies show these capacities decrease in obesogenic/diabetic states and increase with healthy exposures [57–59]; mechanistically, higher β-cell ATP/ADP promotes insulin secretion and lowers glucose [60].

In contrast, three pathways showed an adverse metabolic profile: superpathway of arginine and polyamine biosynthesis (ARG+POLYAMINE-SYN) and superpathway of polyamine biosynthesis I (POLYAMSYN-PWY) showed negative associations: higher pathway abundance correlated with worse liver indices (lower liver sound speed and greater inflammation–fibrosis), higher glucose and CGM variability, and higher body weight/adiposity. Non-oral evidence is directionally consistent—polyamine excess is linked to obesity, inflammation, and impaired insulin signaling, and oral multi-omics work reports elevated oral polyamines in obesity [61, 62]. Ceramide degradation by α-oxidation (PWY-66-388) was positively associated with glycemic deterioration—higher pathway abundance coincided with higher mean glucose and greater CGM variability. This direction is consistent with host evidence: higher ceramide species (core sphingolipids generated in host tissues) are linked to lipotoxicity, insulin resistance, and adverse metabolic outcomes [63, 64].

Pathways with a mixed profile correlated with worse liver health but better glycemia and adiposity. These included the γ-glutamyl cycle (PWY-4041), ADP-L-glycero-β-D-manno-heptose biosynthesis (PWY-0-1241) and the L-methionine superpathway (PWY-5347). These patterns align with host biology: GSH sufficiency improves insulin sensitivity whereas hepatic oxidative stress worsens [65, 66]; mechanistically, ADP-heptose activates ALPK1–NF-κB [67], fitting the liver signal, although prior work reported higher glucose (opposite to our glycemic direction) and there is no evidence linking PWY-0-1241 to body composition. A sulphur-pathway duality (homocysteine vs cystathionine) may underlie PWY-5347 [68]. No prior evidence links (5Z)-dodecenoate biosynthesis II (PWY-7858) to metabolic health.

### 2.4 Metabolic diseases classification

To evaluate the potential of the oral microbiome as a non-invasive diagnostic tool for metabolic diseases, we constructed machine learning models to distinguish between healthy and diseased states across six common metabolic conditions: hypertension, obesity, pre-diabetes, fatty liver, hypercholesterolemia, and cholelithiasis/gallstones. To assess biological relevance while preventing information leakage, we compared two oral-feature model frameworks: (1) a full model incorporating all discovery-stage microbial markers, and (2) a phenotype-filtered model, in which feature selection was restricted to training folds and limited to markers associated with pre-specified clinical phenotypes not used to define case status for the target disease (see Methods 4.8 for selection criteria and leakage controls). For clinical context, we additionally trained baseline classifiers using only age, sex, and BMI under the same cross-validation splits. Biologically informed feature selection consistently and significantly improved classification performance across all six diseases (Fig. 5). For example, in hypertension classification using strain-level features, the phenotype-filtered model increased the AUC from 0.649 to 0.734 (+0.085) and accuracy from 0.672 to 0.741 (+0.069) and out-performed the age/sex/BMI baseline (AUC 0.618; ACC 0.602). Similar performance gains were observed at the pathway level; for example, in classification of Obesity, the AUC improved from 0.627 to 0.728 (+0.101), and Accuracy improved from 0.652 to 0.713 (+0.061). Compared with the age/sex/BMI baseline, oral-feature models achieved higher ACCs and AUCs for five of the six diseases, indicating added classification information beyond simple clinical measures. For obesity, the age/sex/BMI baseline performed similarly or better (AUC 0.775; ACC 0.763), which is expected because BMI is intrinsically related to the obesity definition.

**Fig. 5:**
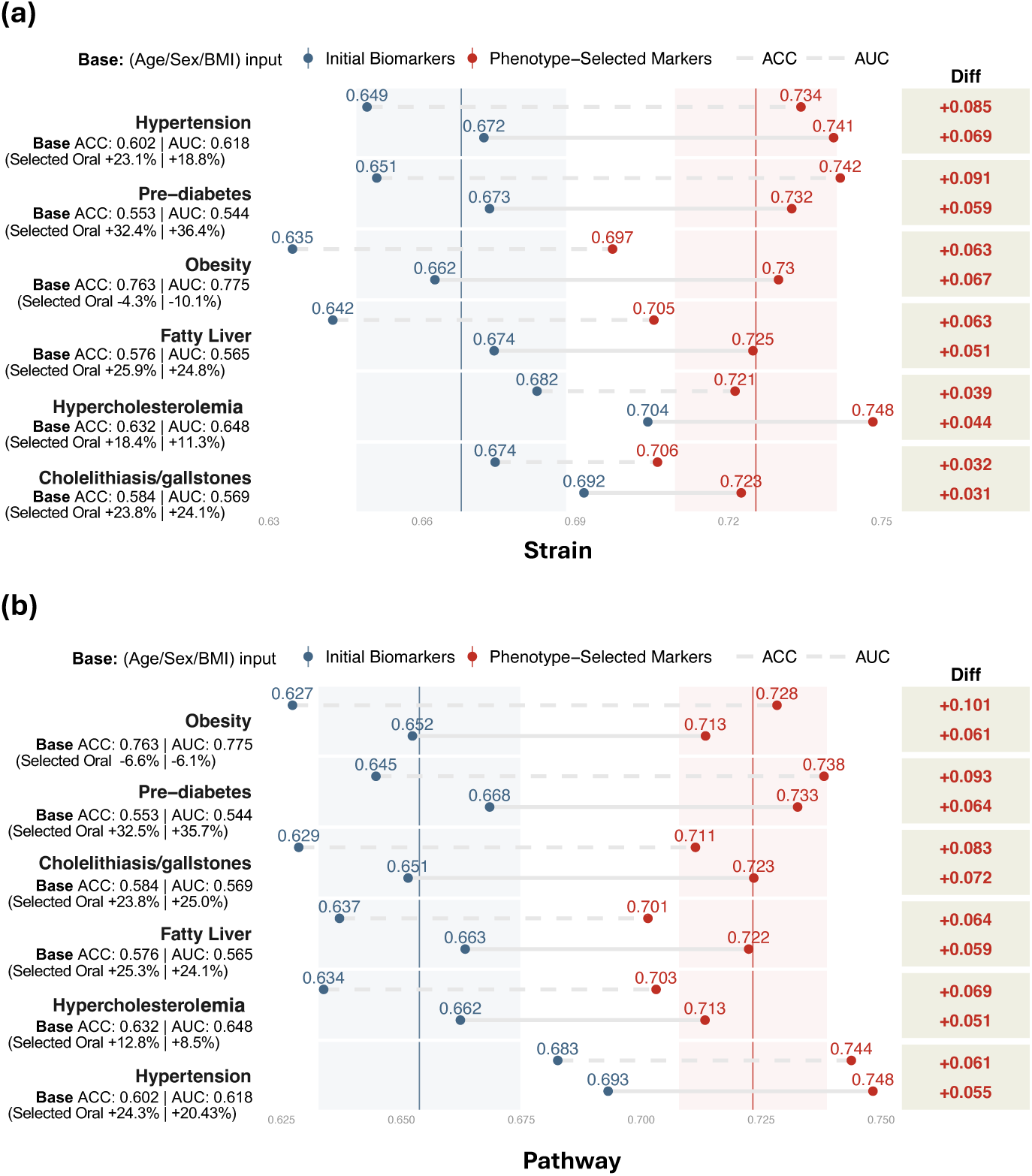
Metabolic disease classification performance using oral microbiome features at two resolutions. (a) Strain level; (b) Pathway level. For each condition, a baseline model using full-feature oral model (blue) is compared with an optimized model restricted to phenotype-selected oral model (red), the selection strategy is detailed in Methods 4.8. For each disease, the left-hand text ‘Base ACC/AUC’ reports the clinical baseline model (age/sex/BMI), together with the relative gain of the phenotype-selected oral model over the baseline. The right-hand Diff column reports the absolute improvement of the optimized model over the baseline for AUC and accuracy (ACC).

To further elucidate the biological signals driving disease classification, we conducted feature importance analyses on the optimized models. Several favorable- and adverse-profile markers previously characterized in Results 2.2 and 2.3 consistently ranked among the most predictive features. In particular, for fatty liver and pre-diabetes, Porphyromonas taxa and the polyamine biosynthesis pathway emerged as high-impact features, whereas the favorable profile strain Corynebacterium durum was strongly associated with healthy status. Overall, results highlight the diagnostic potential of oral microbial features and provide classification-level support of the biological associations identified earlier. In addition, microbial signatures linked to host metabolic phenotypes not only correlate with, but also effectively discriminate metabolic disease states, reinforcing their relevance as potential biomarkers.

### 2.5 Sensitivity analyses accounting for oral health

To assess potential confounding by oral health, we used a brief self-reported questionnaire covering gingival bleeding, gingival abscess/gingivitis, halitosis, and tooth mobility (Table 2). Participants reporting oral antibiotic use within the previous 3 months for gingival/periodontal problems (n=10) were excluded from all oral-microbiome analyses (Methods 4.9).

**Table 2:** Summary of self-reported oral-health indicators in the HPP cohort.

| Category | Description | Count |
| --- | --- | --- |
| **Gingival Bleeding** | Frequent gum bleeding | 544 |
|  | Bleeding “all the time” | 51 |
| **Gingival Abscess/Gingivitis** | Frequent gingival abscess/gingivitis | 95 |
| **Antibiotic Use** | Recent use of oral antibiotics for gingivitis | 10 |
| **Halitosis** | Sometimes noticed by others | 210 |
| **Tooth Mobility** | Slight mobility in a small number of teeth | 547 |
|  | Slight mobility in many teeth | 33 |
|  | Severe mobility | 25 |

First, we repeated the MWAS after excluding participants with pronounced oral problems (n=385), defined as meeting any of the following: gum bleeding reported as “all the time”, gingival abscess/gingivitis reported as frequent, halitosis sometimes noticed by others, or tooth mobility reported as slight mobility in many teeth or severe mobility (Table 2). All other steps (preprocessing, association models, core covariates: age, sex, and smoking status, and Bonferroni correction) were kept identical to the primary analysis. Most Bonferroni-significant associations were retained after exclusion, including 1,702/1,809 strain associations (94.09%), 708,955/843,820 gene family associations (84.02%), and 2,199/2,355 pathway associations (93.38%) (Table S5). Direction concordance was 100% for strains and pathways and 97.37% for gene families. Second, we added two composite oral-health covariates to the main regression models, in addition to age, sex, and smoking status: an inflammation score based on gingival bleeding and gingival abscess/gingivitis, and a periodontal destruction score based on tooth mobility (Methods 4.9). After this adjustment, 1,434/1,809 strain associations (79.27%), 828,984/843,820 gene family associations (98.24%), and 2,308/2,355 pathway associations (98.0%) remained Bonferroni-significant, and all (100%) retained associations had concordant directions of effect (Table S6). We included halitosis only in the exclusion-based definition to keep this sensitivity analysis conservative. We did not incorporate halitosis into the covariate scores because self-reported halitosis is etiologically heterogeneous and subjective [69].

To further assess robustness of key signals, we repeated the most significant oral marker selection procedure using the oral health-adjusted MWAS results. Most original core markers in Results 2.3 were retained (12/15 strains, 13/15 gene families, and 11/13 pathways). The remaining original core markers were not re-selected as the top features after oral health adjustment, but they remained Bonferroni significant for at least one phenotype, indicating that oral-health adjustment primarily affected marker ranking rather than eliminating the underlying associations. Finally, for metabolic disease classification, we additionally removed participants meeting the same pronounced oral problem definition and refit the same classification pipeline on the reduced case sets with matched controls. Model performance changed modestly after excluding cases with pronounced oral problems (Table S7). Across diseases, AUC shifts ranged from −0.031 to +0.015, with the largest decreases observed for pre-diabetes (−0.031) and obesity (−0.027) in strain-based models and for fatty liver (−0.020) in pathway-based models, while cholelithiasis/gallstones showed a small increase (+0.015).These results suggest that the oral-feature classifiers retain discernible signal and are not driven solely by participants with pronounced self-reported oral problems.

### 2.6 Validation in an independent cohort

To ensure the reproducibility and generalizability of our findings, we conducted a validation analysis in National Health and Nutrition Examination Survey (NHANES) Oral Microbiome project [70], NHANES is an independent large-scale cohort comprising 20,293 individuals. This cohort included genus-level oral microbiome profiles paired with matched phenotypic data (*n* = 10,081 males, 10,212 females; age range: 1–80 years; mean ± SD: 32.02 ± 24.75 years; BMI: 25.64 ± 7.73 kg/m^2^; waist circumference: 87.10 ± 22.31 cm). To ensure methodological comparability, metagenomic processing, phenotype definitions, and association modeling were performed identically to those applied in the HPP discovery cohort (Methods 4.10). Because the replication cohort provides only genus-level profiles and a limited set of overlapping phenotypes, validation focused on directional replication of the HPP BMI- and waist-circumference associations at the genus level.

We assessed the external reproducibility of the HPP strain-level findings by testing their genus-level counterparts in an independent cohort. Because strain-level resolution was not available in the replication dataset, the BMI-associated strains were mapped to genus and collapsed to 28 non-redundant genera via a many-to-one mapping procedure (Methods 4.10). Among these 28 genera, 11 (39.3%) were significant with the same effect direction, and 8 of these (72.7%) demonstrated concordant effect estimates. Similarly, for waist circumference–associated strains, 25 corresponding genera were evaluated; 9 (36.0%) were replicated, and 8 (88.9%) displayed consistent effect directions. To complement the strain-to-genus validation, we performed a parallel genus-level replication analysis. Among the 18 BMI-associated genera in the discovery cohort, 10 (55.6%) were also significantly associated in the replication cohort, with 9 (90.0%) showing directional concordance. For waist circumference, 8 of the 16 associated genera (50.0%) were replicated, 8 (100.0%) of which exhibited consistent directions of effect.

In conclusion, these multi-level replication analyses demonstrate partial but directionally consistent reproducibility under genus-level resolution, thereby supporting the robustness and external validity of the key oral microbial associations identified in the HPP cohort.

## 3 Discussion

In recent years, the oral microbiome has been consistently linked to host metabolic health, with associations reported for obesity, type 2 diabetes, and non-alcoholic fatty liver disease (NAFLD) [9, 10]. Therapeutic approaches targeting the oral ecosystem, such as periodontal treatment, oral probiotics, and microbiota modulators, have shown potential to improve liver function, glycemic control, and body fat composition, and are increasingly regarded as adjunctive treatments for metabolic disorders [71–73].

Despite recent advances, systematic, population-scale evidence linking the oral microbiome at strain-level and functional (gene family and pathway) resolution to metabolic health remains limited. To address this gap, we analyzed 9,431 participants in the Human Phenotype Project and linked oral metagenomes to 44 phenotypes across liver ultrasonography, CGM, and body composition. We identified 213 strains, 124,603 gene families, and 299 pathways that were associated with at least one phenotype. Associations were pleiotropic yet system-specific: most features spanned multiple phenotypes, but about half were confined to a single system. Strain signals concentrated in body composition, with strong positive links to adiposity and weaker links to lean mass. In the functional layer, CGM contributed most associations and system-specific signals, with positive directions for mean glucose and hyperglycemia risk and negative directions for hypoglycemia risk and glycemic variability. The liver domain showed the fewest associations and the directions of the strains and pathways differed between inflammation and fibrosis.

Next, we prioritized features with the broadest and most robust relationships to metabolic health. Where strain-level reports were unavailable in the literature, we contextualized findings at the species or genus level and, when oral precedents were limited, with non-oral evidence anchored by relevant metabolites and pathway products. With few exceptions, directions were concordant with prior reports. At the strain layer, beneficial strain profiles aligned with oxygen tolerance or nitrate use and with lower liver inflammation and fibrosis. Detrimental profiles were enriched for anaerobic, inflammophilic fermenters associated with higher glucose, steatosis, and visceral adiposity. Mixed profiles showed system-dependent trade-offs. At the gene-family layer, beneficial signals clustered in core biosynthesis, while detrimental signals concentrated in translation and replication modules and in glutamine-dependent NAD^+^ synthetase. These patterns point to partial uncoupling between liver status and glucose control. At the pathway layer, UDP-N-acetylglucosamine and UDP-N-acetylgalactosamine biosynthesis, L-valine biosynthesis, and de novo purine biosynthesis aligned with healthier liver indices, more stable CGM profiles, and favorable fat distribution. In contrast, polyamine biosynthesis and ceramide alpha-oxidation aligned with adverse outcomes.

We also trained metabolic disease classification models using oral features that were significantly associated with metabolic phenotypes. Models built from selected features outperformed full feature models on several conditions (detailed results shown in Results 2.4), indicating stronger and more coherent links to disease status.

This study advances understanding of oral–metabolic links but has limitations. First, participants were recruited from a single geography in Israel, which may limit generalizability, and replication was feasible only at the genus level because strain-level data were unavailable in the external cohort, leading to weaker effects than at the strain level despite careful matching. Second, our oral microbiome profiles were derived from standardized bilateral buccal-mucosa swabs and therefore reflect a single oral niche rather than the whole mouth microbiota. In addition, we did not systematically record the sampling time of day or very short-term behaviors (e.g., recent speaking, or salivary-flow changes), and oral health status was assessed by brief self-report rather than comprehensive clinical dental phenotyping; together, these factors may add variability and leave residual confounding. Third, functional inference relies on metagenomic annotation and reference databases; low abundance, rare, or poorly characterized taxa and gene families may be underestimated. Fourth, the present analyses are cross-sectional; they provide associations and generate hypotheses but do not establish causality.

Despite these limitations, high-resolution strain and functional evidence remains limited in the field, and our cohort-scale map directly addresses this gap. The sample size and breadth of phenotyping support robustness and potential transportability relative to small cohorts. As an observational resource, our association map and the prioritized oral markers can provide reusable candidates for future longitudinal and mechanistic studies to test causality. From an applied perspective, oral markers merit evaluation as noninvasive tools for metabolic risk stratification and early screening. On the other hand, reinforcing beneficial community functions and selectively suppressing detrimental pathways could guide the development of microbiome-based therapeutics for metabolic disease.

## 4 Methods

This research complies with all relevant ethical regulations. The Human Phenotype Project (HPP) was conducted in accordance with the principles of the Declaration of Helsinki and was approved by the Weizmann Institutional Review Board (IRB) at the Weizmann Institute of Science (reference no. 1719-1). All participants provided written informed consent. At HPP registration, participants were asked to report their date of birth and sex; therefore, sex was self-reported, and no separate gender variable was used in this study.

### 4.1 Description of HPP cohort

The Human Phenotype Project (HPP) is a large-scale, longitudinal deep-phenotyping cohort of 9,431 individuals aged from 38 to 72. Eligibility criteria: HPP recruits community-dwelling volunteers from Israel with the aim of enrolling generally healthy adults. Key exclusion criteria at recruitment included pregnancy or ongoing fertility treatments, frequent hospitalizations (e.g., *>*3 in the previous year), unintentional weight loss (*>*5% in the previous year), prior or active malignancy, and major chronic or unstable cardiovascular, neurologic/psychiatric, respiratory, renal, or gastrointestinal conditions (including inflammatory bowel disease and liver cirrhosis). Since January 2019, the HPP has concurrently collected a wide range of multimodal data covering multiple major physiological systems. These data modalities include: tabular data recording personal medical history and physiological information; deep molecular data encompassing genomics, proteomics, and microbiomics; time-series data reflecting dynamic physiological changes, such as continuous glucose monitoring; and key imaging data, including liver ultrasound and DXA scans.

In this study, we used metagenomic data from buccal swabs in the HPP cohort and combined these with three core metabolic assessments: (1) liver health status, obtained by liver ultrasonography; (2) dynamic glucose homeostasis, acquired through CGM; and (3) body composition, including visceral adipose tissue, as measured by DXA. Participants reporting recent oral antibiotic use (*<*3 months) for gingival or periodontal problems (n = 10) were excluded from the oral-microbiome analyses in this study. Baseline questionnaires recored self-reported physician diagnosed metabolic conditions (used as case labels in Results 2.4) and self-reported oral health (used for sensitivity analyses in Results 2.5). This is consistent with the HPP recruitment strategy, which enriches for generally healthy adults by excluding major acute or unstable systemic diseases, while allowing common stable conditions that are captured in the baseline questionnaires. These data provide a basis for analyzing the potential relationships between the oral microbiome and metabolic phenotypes.

### 4.2 Oral Microbiome Collection and Preprocessing

Oral microbiome samples were collected via standardized bilateral buccal-mucosa swabs during the baseline visit following a unified sampling SOP (Table S1) and processed at Gencove laboratories for DNA extraction and library preparation. Buccal mucosa is a standard oral sampling site used in large cohort studies (e.g., the Human Microbiome Project [74]), and our protocol follows published best-practice recommendations for oral-microbiome sampling [75]. Briefly, trained staff wearing sterile gloves used single-use flocked swabs to rub and rotate along the left and right buccal mucosa with moderate pressure, while deliberately avoiding contact with teeth, gingival crevices, tongue dorsum, and saliva pools, to minimize contamination from supragingival plaque and transient saliva. Participants were asked to refrain from tooth-brushing or antiseptic mouthwash for ≥4 h and from eating for ≥1 h (water allowed). Immediately before sampling, they gently rinsed once with water and waited 10–15 min. Swabs were placed into tubes immediately, frozen at −80 ^◦^C within 2 h of collection, and shipped on dry ice for downstream sequencing. Accordingly, these samples therefore primarily reflect the buccal mucosal community rather than a whole-mouth or saliva composite [76, 77]. Subsequent whole genome sequencing was performed at Neogen GeneSeek Laboratories on an Illumina NovaSeq platform, utilizing a 150-bp paired-end read configuration.

Raw sequencing reads were first aligned to the human reference genome to isolate non-human reads. These reads were then subjected to quality control using Trimmomatic to remove sequencing adapters and low-quality sequences. The quality-controlled reads were then processed through a dual analysis pipeline. The taxonomic composition was profiled with MetaPhlAn 4 to determine the relative abundance of microbial taxa from kingdom to strain level. Concurrently, functional potential was characterized with HUMAnN 3.6 to quantify the abundance of microbial gene families and metabolic pathways. The resulting taxonomic and functional profiles, derived from the processed non-aligned (non-human) reads, were preprocessed for microbiome analyzes.

After host-depletion and read-level QC, we retained features (strains, gene families, pathways) with ≥ 200 non-zero observations across samples. For each retained feature, we defined a feature-specific detection threshold as the 1st percentile of its positive abundances and replaced zero entries with half of this threshold to avoid large pseudo-count bias.

Let 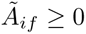 denote the resulting (zero-replaced) abundance of feature *f* in sample *i*, and let 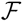 be the feature set. We then performed within-sample total-sum normalization and scaling. To stabilize variance and improve approximate normality in downstream association models, we subsequently applied a base-10 log transformation. These steps are defined as follows:

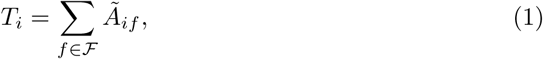

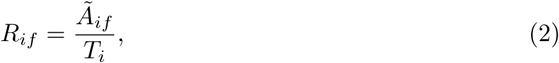

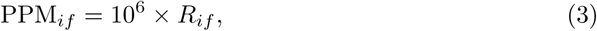

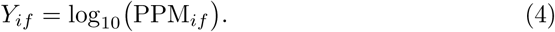

Because of zero replacement, PPM*_if_ >* 0 and the logarithm is well-defined. All steps were applied identically to strains, gene families and pathways.

We further filtered the taxonomic attributions for each pathway provided by HUMAnN. To reduce noise from low-prevalence organisms, only genera and species that contributed to a pathway’s abundance in at least 200 samples were retained for subsequent analysis.

### 4.3 Metabolic health phenotypes collection and preprocessing

In this study, we selected phenotypes related to liver health, glucose homeostasis, and body composition, as these are well-established and critical indicators of metabolic health. The detailed descriptions are listed as follows:

#### Liver

Participant liver health is quantitatively assessed using a Supersonic Aixplorer MACH 30 ultrasound system (Hologic, USA) with a C6-1X convex transducer. Following a standardized protocol, measurements are taken from three distinct locations within the right liver lobe via an intercostal approach while participants are supine and holding their breath. To ensure measurement quality for elasticity and viscosity, a stability index greater than 90% was required. This procedure yields key quantitative metrics for assessing fibrosis (Elasticity), inflammation (Viscosity), and steatosis (Sound Speed and Attenuation), providing a final dataset with values in units such as kPa and Pa·s.

#### CGM

Dynamic glucose homeostasis is assessed using the FreeStyle Libre ProIQ Flash continuous glucose monitoring (CGM) device. During the baseline visit, a sensor is placed on the back of the participant’s upper arm and activated for a two-week period, recording interstitial glucose levels every 15 minutes. Concurrently, participants log information on their medication, physical activity, and sleep hours using a dedicated application (10K application). This method yields a high-resolution time-series of glucose levels, from which various summary metrics of glycemic control and variability (e.g., mean glucose, coefficient of variation, time in range) are calculated using the iglu R package [78].

#### Body Composition

Body composition and bone mineral density are assessed by Dual-energy X-ray Absorptiometry (DXA) using a GE Lunar Prodigy Advance device (GE Healthcare, USA). For the total body scan, participants are positioned supine after removing all metal items and heavy clothing. This scan provides detailed measurements of fat mass, lean mass, and percent body fat for the whole body and for specific regions (arms, legs, trunk, android, and gynoid). Furthermore, the GE CoreScan software is used to quantify Visceral Adipose Tissue (VAT) and Subcutaneous Adipose Tissue (SAT) within the android region, while bone mineral density is specifically measured at the bilateral femur necks and the L1-L4 lumbar spine.

For each phenotype *p* from the liver, CGM, and body composition systems, we performed the data cleaning process. To obtain robust location and scale, we computed the shortest interval covering 95% of the non-missing observations and estimated the mean *µ_p_*and standard deviation *σ_p_* within that interval. We then removed extreme outliers defined by the condition |*x_ip_* −*µ_p_*| *>* 8*σ_p_*. For the remaining values, we applied winsorization at ±5*σ_p_*:

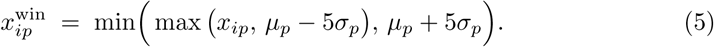

These cleaned phenotypes were used in all downstream microbiome–phenotype association analyses.

### 4.4 Identification of significant strain, gene family, pathway associations

Associations between oral strain, gene family, and pathway profiles and 44 host metabolic phenotypes were assessed using ordinary least squares (OLS) linear regression models. These models were implemented with the statsmodels package (v0.14.4) in Python (v3.11), with adjustments for age, sex, and smoking status. Age, sex, and smoking status were included as covariates because large population-based studies have identified them as major host determinants of oral microbial diversity and composition [70, 79]. This minimal, unified covariate set helps reduce confounding while keeping the 44 phenotype-specific models directly comparable. The analysis environment was Ubuntu 22.04, utilizing the NumPy (v1.26) and pandas (v2.2) libraries. The microbial features were first preprocessed using Methods 4.2, while phenotypic variables were cleaned using Methods 4.3. For each association, we estimated effect coefficient and corresponding P-values. Bonferroni correction was applied separately to each feature layer. For layer *t* ∈ {strain, gene, pathway}, with *N_t_* tested features and *K* = 44 phenotypes, the per-test significance threshold was set at *p <* 0.05*/*(*N_t_* × *K*).

### 4.5 Mining for core gene family

Given the large number of gene families identified (N = 355,674), we anticipated significant challenges from high inter-correlation and functional redundancy. To address this, we designed and implemented two orthogonal approaches: a pre-association decorrelation filter and a post-association independence pruning step. The pre-association filter was applied prior to association testing to generate a reduced set of representative, uncorrelated gene families. In contrast, the post-association pruning was applied after the main analysis to distill the list of significant associations down to a set of statistically independent signals for each phenotype.

#### Pre-association decorrelation filter

To mitigate collinearity among bacterial gene families prior to association testing, we applied a pre-association decorrelation filter. All gene families were randomly partitioned into batches of up to 3,000 features. Within each batch, we computed pairwise Spearman rank correlations and applied average-linkage hierarchical clustering using a correlation cutoff of 0.3 to define clusters. For each cluster, we generated a candidate subset by ranking features according to their mean abundance across samples and retaining the top 5%. Pairwise proximity among candidates was quantified as

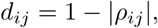

and the cluster representative was defined as the feature with the smallest mean distance to all other candidates.

After identifying representative gene families from each batch, we pooled these features and reapplied the clustering procedure iteratively for three successive rounds to further suppress inter-feature redundancy. This iterative decorrelation process produced a final feature set with minimal mutual correlation. The resulting retained gene families exhibited an average pairwise correlation of 0.01 (mean absolute Spearman’s *ρ*, *P <* 1 × 10^−3^) with an overall standard deviation of 0.26, indicating substantial independence among the selected features.

#### Post-association pruning

After identifying gene families associated with metabolic health, we applied a pruning strategy to determine a subset of gene families exhibiting independent associations with the relevant host metabolic phenotypes. After the association analysis (Methods 4.4), for each metabolic phenotype *Y_k_* we retained gene families *g* with Bonferroni-adjusted p-values 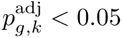 and their regression coefficients 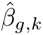, where *X_g_*denotes the abundance of gene family *g*. To place coefficients on a phenotype-specific, unit-free scale, we computed the standardized effect size

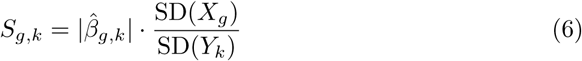

Within phenotype *k*, candidates were ranked by a joint rank that treats significance and effect size symmetrically:

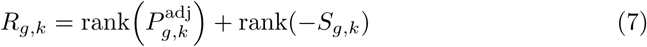

For each phenotype *Y_k_*, we selected the top-ranked candidate (by *R_g,k_*), and added it to the approximately independent set, and removed any remaining candidate whose Spearman correlation with the lead satisfied *ρ* ≥ 0.30. We iterated until no candidates remained, yielding a set of independent gene families for each phenotype.

### 4.6 Identification of key oral microbial features associated with metabolic health

We developed a systematic selection strategy to identify the key oral microbial strains, gene families, and pathways most strongly and broadly associated with host metabolic phenotypes. To capture system-level diversity, all phenotypes were first organized into three metabolic domains: liver, continuous glucose monitoring (CGM), and body composition. Within each system, we quantified the association breadth of every microbial feature by counting the number of phenotypes with which it showed a significant relationship (Bonferroni-corrected *p <* 0.05). Strains and pathways were then ranked according to this association count, such that features linked to a larger number of phenotypes received higher priority. For gene families, we began with the post-association pruned set obtained after correlation-based clumping (Methods 4.5), ensuring that the candidate features were functionally independent and not phenotypically redundant. These pruned gene families were then ranked within each system using the same association-based metric. From each of the three systems, we selected the top five ranked features per data type (strain, gene family, and pathway) as the key microbial components for downstream analyses. In order to avoid the repeated representation of cross-system features, we performed de-duplication step by merging identical features across systems into a unified, non-redundant set. This hierarchical selection procedure prioritized features that were not only statistically significant but also robustly associated across multiple related phenotypes, resulting concise and biologically meaningful set of core oral microbial markers.

### 4.7 Grouping of oral microbial features

Each microbial feature (strain, gene family, or pathway) was grouped into a favourable metabolic profile, adverse metabolic profile, or mixed profile by integrating its association directions across physiological systems (liver, CGM, and body composition), without cross-system averaging. These profile labels are defined solely by the direction of association after phenotype orientation and do not imply causality. Starting from Bonferroni-significant feature–phenotype associations, we oriented all phenotypes to a common health axis using a pre-specified mapping: for phenotypes where higher values indicate better health, the original association sign was retained; for those where lower values indicate better health, the sign was inverted. The full phenotype-specific orientation scheme is provided in Supplementary Table S3. This approach to direction-of-effect synthesis, coupled with transparent reporting of grouping and weighting rules, follows established best practices for evidence synthesis without meta-analysis.

Within each system, phenotypes were grouped into pre-defined clinical domains, and all significant associations within a given system–domain combination were collapsed into a single domain-level vote using a weighted-majority scheme. This procedure preserved directional coherence while reducing redundancy among correlated phenotypes. Association-level weights preserved the dynamic range of evidence but capped extremes, *w* = min{3, − log_10_(*P*_bonferroni_)}; domain weights were normalized so the sum per system equaled 1. For system *s*, we defined the weighted positive proportion below:

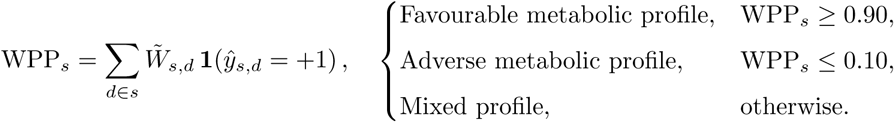

where 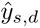 denotes the domain-level vote and 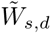 the normalized domain weight. These grouping follow the same three-system framework and domain definitions (described in the main text).

### 4.8 Metabolic diseases classification

We modeled each condition independently in the HPP cohort using balanced case– control data. Disease labels were derived from self-reported physician-diagnosed conditions. Case counts were: Hypercholesterolemia (*n* = 2409), hypertension (*n* = 695), pre-diabetes (*n* = 449), obesity (*n* = 589), fatty liver (*n* = 623), and cholelithiasis/gallstones (*n* = 650). For each disease, an equal number of controls was randomly sampled from the HPP healthy pool; sampling was performed separately for each disease among participants carrying an explicit healthy label. Control sampling was performed once per disease to construct a fixed balanced dataset, which was then used across all cross-validation folds. Models were fit at two feature layers—strain and pathway—from the corresponding oral microbiome abundance profiles. We trained the LightGBM classifier and evaluated performance using stratified five-fold cross-validation (80% training / 20% testing per fold). No separate validation set was used, and AUC/ACC values were averaged across the five test folds. To benchmark against established clinical risk factors and quantify the added value of oral features, we additionally trained an age/sex/BMI baseline classifier on the same balanced dataset under identical cross-validation splits.

Within each training fold, candidate clinical phenotypes were pooled from liver ultrasound, CGM, and body composition phenotypes, and self-reported metabolic diseases. For the target disease label (case vs control), we ran univariate ANOVA F-tests for each candidate phenotype using SelectKBest and f classif from scikit-learn (V1.7.1), obtaining an F score and P value per phenotype, where the F statistic quantifies how much the mean value of that phenotype differs between cases and controls relative to the variability within each group (a larger F means better separation). Phenotypes were ranked by F score (ties broken by P value), and we retained the top 20 ranked phenotypes as a simple, transparent filter that highlights phenotypes most different between cases and controls while keeping this screening step compact and conservative. These phenotypes are then used only to select informative oral markers; they themselves are not included as predictors in the final microbiome-based classifiers. Using training-fold data only, we identified oral markers (strains or pathways) that met Bonferroni-adjusted P value *<* 0.05 for any of the top 20 phenotypes, using the criteria defined in Results 2.2. We then trained the classifier on this training-defined marker set. In the test fold, the top-20 list and marker set were fixed in advance and applied without recomputation or reselection to prevent data leakage. For each disease and feature layer, performance for the phenotype-filtered and full-feature pipelines was summarized across the same five cross-validation splits to enable paired comparisons.

### 4.9 Sensitivity analyses accounting for oral health

To evaluate potential confounding by oral health status in oral–metabolic phenotype associations, we conducted oral-health sensitivity analyses using a brief self-reported oral-health questionnaire covering gingival bleeding, gingival abscess/gingivitis, halitosis, and tooth mobility (summary statistics in Table 2). Participants reporting oral antibiotic use within the previous 3 months for gingival or periodontal problems (n=10) were excluded from all oral-microbiome analyses.

First, we repeated the association analysis after excluding participants with pronounced oral problems, defined as any of the following: gum bleeding reported as all the time; gingival abscess/gingivitis reported as frequent; halitosis sometimes noticed by others; or tooth mobility reported as slight mobility in many teeth or severe mobility, in total (n=385). All other steps (preprocessing, OLS models, core covariates: age, sex, and smoking status, and Bonferroni correction) were kept identical to the primary analysis in Methods 4.4. For metabolic disease classification, we additionally removed participants meeting the same pronounced oral problem definition in case and control (excluded/total cases: fatty liver 34/623, hypercholesterolemia 103/2409, hypertension 34/695, pre-diabetes 31/449, obesity 42/589, and cholelithiasis/gallstones 16/650) and refit the same classification pipeline on the reduced case sets with matched controls.

Second, we added two composite oral-health covariates (an inflammation score and a periodontal destruction score) to the main regression models, alongside age, sex, and smoking status, to account for residual oral-health variation. Previous studies support self-reported gum bleeding (including bleeding on brushing) as an indicator of gingival inflammation and self-reported tooth mobility as an indicator of periodontitis-related tissue breakdown; based on this evidence, we derived these two composite oral-health covariates for sensitivity analyses [80–82]. Details are as follows: **(i)** Inflammation Score (0–2) derived from self-reported gingival bleeding and gingival abscess/gingivitis: 0, neither frequent gum bleeding nor frequent abscess/gingivitis; 1, either (frequent gum bleeding, but not “all the time”, without frequent abscess/gingivitis) or (frequent abscess/gingivitis without frequent gum bleeding); 2, gum bleeding reported as “all the time” or both frequent gum bleeding and frequent abscess/gingivitis. **(ii)** Periodontal Destruction Score (0–3) derived from tooth mobility (0: none; 1: slight mobility in a small number of teeth; 2: slight mobility in many teeth; 3: severe mobility). All other steps were kept identical to the primary analysis.

Halitosis was included only in the exclusion-based definition of pronounced oral problems to make this sensitivity analysis conservative, but was not incorporated into the covariate scores because self-reported halitosis is heterogeneous and subjective with limited agreement with objective malodor assessments [69], and including it as a covariate could introduce measurement noise.

### 4.10 Replication study

We evaluated external reproducibility in the U.S. National Health and Nutrition Examination Survey (NHANES) Oral Microbiome project [70], an independent cohort comprises 20,293 participants with oral microbiome profiles at the genus level and matched anthropometrics (females: *n* = 10,212; males: *n* = 10,081; age 32.02 ± 24.75 years; BMI 25.64 ± 7.73,kg/m²; waist circumference 87.10 ± 22.31cm). This validation dataset combines the 2009–2010 and 2011–2012 NHANES cycles. Oral-rinse samples were profiled by 16S rRNA V4 (515F/806R) sequencing on Illumina HiSeq 2500 and processed by the U.S. Centers for Disease Control and Prevention into amplicon sequence variant (ASV)-based relative-abundance tables summarized at the phylum-to-genus levels. Anthropometric and clinical covariates were obtained from the corresponding continuous NHANES data files, and associated sequencing QC data are available under BioProject accession PRJNA896165. To harmonize analyses with the discovery cohort, we applied the same preprocessing and association framework where applicable (Methods 4.2, 4.3, and 4.4).

Because the replication resource does not provide strain resolution or functional profiles, we assessed robustness of strain–phenotype signals at the genus level using two complementary strategies. Strategy 1 (strain→genus mapping): each discovery strain significantly associated with BMI or waist circumference was mapped to its corresponding genus (many–to–one). Replication was declared for a genus if it was significantly associated with the same phenotype in NHANES and the effect direction matched that observed in discovery. Strategy 2 (genus–level re-analysis in discovery): to mitigate resolution mismatch, we recomputed genus–phenotype associations within the discovery cohort using the same models and thresholds, then tested whether the corresponding genera in NHANES showed significant effects in the same direction. Missing phenotypes or covariates were handled by listwise exclusion.

No replication was attempted for gene families or pathways because such functional abundances are unavailable in NHANES oral data.

## 5 HPP data availability

HPP data used in this study are available under controlled access within the HPP Trusted Research Environment (TRE). Access is restricted because oral shotgun metagenomic data were generated from buccal-swab samples, raw sequencing files contain human-derived reads, the host-depleted files may retain residual host-derived sequences, and the linked individual-level clinical phenotype data are privacy-sensitive. Bona fide researchers may request access for in-place analysis within the TRE by contacting; further information is provided on the HPP data access website. Requests are reviewed under the applicable HPP data-use agreement and IRB framework and are typically reviewed within a few days. If approved, access is provided within the TRE. For the NHANES replication analyses, oral microbiome relative abundance data and BMI, waist circumference data are obtained from NHANES website. Access to NHANES raw oral microbiome sequencing data at the participant level must be requested through the NCHS Research Data Center (RDC) application process.

## 6 Code availability

Code used in this study is available at GitHub: https://github.com/haochen-MBZUAI/HPP-Oral-Microbiome.

---

**Table S1:** Summary of the buccal-swab sampling SOP.

| Stage | Key Actions | Purpose |
| --- | --- | --- |
| **Pre-sampling Restrictions** | No tooth-brushing or antiseptic mouthwash for $\geq 4$ h<br>Fasting for $\geq 1$ h (small amounts of water allowed) | To minimize acute perturbations from recent hygiene procedures and food intake. |
| **On-site Preparation** | Gently rinse mouth once with a small volume of water<br>Wait 10–15 min before sampling | To remove visible food debris while minimally disturbing the resident microbial community. |
| **Sampling Tools** | Sterile gloves worn by study nurses<br>Single-use flocked swabs | To ensure aseptic technique and maximize sample collection efficiency. |
| **Sampling Procedure** | Rub and rotate swab along left and right buccal mucosa<br>Apply moderate pressure | To effectively enrich for mucosa-associated microbial communities. |
| **Critical Avoidance** | Explicitly avoid contact with:<br>Teeth and gingival crevices<br>Tongue dorsum<br>Saliva pools | To reduce contamination from plaque or pooled saliva bacteria (improving tissue specificity). |
| **Storage & Transport** | Swabs placed in tubes immediately<br>Frozen at $-80^{\circ}\text{C}$ within 2 h<br>Shipped on dry ice | To prevent DNA degradation and preserve the microbiome profile for sequencing. |

**Table S2:** Extended summary statistics of all metabolic phenotypes analyzed in this study. Values are mean (s.d.), organized by liver ultrasound phenotypes, CGM-derived phenotypes, and body composition phenotypes.

| Phenotype | Male<br>n=4474(47.4%) | Female<br>n=4957(52.6%) | All<br>n=9431 |
| --- | --- | --- | --- |
| **Liver Health** |  |  |  |
| **Liver Steatosis** |  |  |  |
| Liver sound speed (m/s) | 1547.36 (23.07) | 1539.51 (23.69) | 1543.31 (23.72) |
| Liver attenuation (dB/cm/MHz) | 0.41 (0.10) | 0.41 (0.10) | 0.41 (0.10) |
| **Fibrosis** |  |  |  |
| Liver Elasticity (kPa) | 5.43 (1.25) | 4.68 (0.94) | 5.04 (1.17) |
| Shear Wave Velocity (m/s) | 1.34 (0.14) | 1.24 (0.12) | 1.29 (0.14) |
| **Inflammation** |  |  |  |
| Liver Viscosity (Pa·s) | 1.70 (0.24) | 1.62 (0.27) | 1.66 (0.26) |
| Shear Wave Dispersion ((m/s)/kHz) | 2.46 (0.45) | 2.32 (0.50) | 2.39 (0.48) |
| **Glycemic Control (CGM)** |  |  |  |
| **Average Glucose Level** |  |  |  |
| Mean Glucose (mg/dL) | 97.35 (13.74) | 94.08 (11.23) | 95.63 (12.59) |
| Median Glucose (mg/dL) | 94.31 (11.72) | 91.30 (10.67) | 92.73 (11.28) |
| Glucose Management Indicator (GMI, %) | 5.64 (0.33) | 5.56 (0.27) | 5.60 (0.30) |
| **Glycemic Variability** |  |  |  |
| Glucose IQR (mg/dL) | 17.98 (6.78) | 17.02 (4.98) | 17.48 (5.92) |
| Coefficient of Variation (CV, %) | 15.95 (4.18) | 16.15 (4.00) | 16.05 (4.09) |
| Mean Absolute Deviation (MAD, mg/dL) | 12.62 (4.69) | 11.91 (3.38) | 12.25 (4.05) |
| Mean of Daily Differences (MODD, mg/dL) | 13.62 (4.93) | 12.88 (4.04) | 13.23 (4.50) |
| Mean Absolute Glucose (MAG) | 10.85 (2.94) | 10.64 (2.73) | 10.74 (2.83) |
| SD of ROC | 0.4430 (0.1002) | 0.4352 (0.0944) | 0.4389 (0.0973) |
| SD between Time Points (Time-of-Day SD) | 7.73 (2.98) | 7.97 (2.82) | 7.86 (2.91) |
| SD between Days (Interday SD) | 13.81 (3.76) | 13.70 (3.53) | 13.75 (3.64) |
| Within-Hour SD | 5.29 (1.24) | 5.18 (1.16) | 5.23 (1.20) |
| **Hyper/Hypoglycemia Risk** |  |  |  |
| High Blood Glucose Index (HBGI) | 0.23 (0.94) | 0.14 (0.36) | 0.18 (0.70) |
| Low Blood Glucose Index (LBGI) | 2.18 (1.91) | 2.57 (1.94) | 2.39 (1.93) |
| Time > 140 mg/dL (%) | 2.44 (3.15) | 2.05 (2.83) | 2.23 (3.00) |
| Time > 180 mg/dL (%) | 0.08 (0.19) | 0.07 (0.18) | 0.07 (0.19) |
| Average Daily Risk Range (ADRR) | 10.32 (4.70) | 11.27 (4.78) | 10.82 (4.77) |
| **Glycemic Control Quality** |  |  |  |
| Composite Glycemic Index (COGI) | 0.89 (0.16) | 0.86 (0.17) | 0.87 (0.16) |
| Diabetes Risk Assessment (GRADE) | 1.37 (1.23) | 1.24 (0.85) | 1.30 (1.05) |
| GRADE Euglycemia | 68.08 (21.92) | 65.30 (21.69) | 66.62 (21.84) |
| **Body Composition** |  |  |  |
| **Overall Adiposity** |  |  |  |
| BMI (kg/m<sup>2</sup>) | 26.70 (3.77) | 25.75 (4.40) | 26.21 (4.13) |
| Total Fat Mass (kg) | 23.16 (8.05) | 25.98 (8.56) | 24.60 (8.43) |
| Total Body Fat (%) | 28.52 (6.32) | 38.71 (6.79) | 33.71 (8.31) |
| Total Mass (kg) | 79.48 (12.53) | 65.66 (11.89) | 72.44 (14.03) |
| **Fat Distribution** |  |  |  |
| Waist Circumference (cm) | 94.62 (10.87) | 84.07 (10.99) | 89.24 (12.14) |
| Hip Circumference (cm) | 102.43 (7.43) | 102.57 (9.37) | 102.51 (8.48) |
| Waist-to-Hip Ratio (WHR) | 0.92 (0.07) | 0.82 (0.07) | 0.87 (0.09) |
| Android Fat Mass (g) | 2327.65 (1105.68) | 2071.05 (963.79) | 2196.94 (1043.69) |
| Android Fat % | 0.35 (0.10) | 0.40 (0.10) | 0.37 (0.10) |
| VAT Mass (g) | 1211.70 (748.66) | 647.77 (458.77) | 924.62 (679.53) |
| VAT Volume (cm<sup>3</sup>) | 1279.13 (776.47) | 686.63 (486.30) | 977.44 (709.95) |
| Trunk Fat Mass (g) | 13370.60 (5358.01) | 12929.46 (5013.76) | 13145.68 (5189.87) |
| Trunk Fat % | 0.32 (0.08) | 0.38 (0.09) | 0.35 (0.09) |
| SAT Mass (g) | 1397.79 (683.01) | 1600.33 (749.12) | 1500.89 (724.50) |
| SAT Volume (cm<sup>3</sup>) | 1481.66 (723.99) | 1696.35 (794.07) | 1590.95 (767.97) |
| **Lean Mass** |  |  |  |
| Total Lean Mass (kg) | 56.33 (6.82) | 39.67 (4.98) | 47.84 (10.23) |
| Android Lean Mass (kg) | 4.07 (0.57) | 2.90 (0.43) | 3.48 (0.77) |
| Trunk Lean Mass (kg) | 27.10 (3.31) | 19.63 (2.48) | 23.29 (4.74) |

**Table S3:** Description of All Metabolic Health Phenotypes. The “Health direction” column defines the sign convention used to align phenotypes to a common health axis (Higher = higher is healthier; Lower = lower is healthier; “-” = not oriented); this is for direction harmonization only and does not imply causality.

| Phenotype | Short Description | Health direction |
| --- | --- | --- |
| **Liver Health** |  |  |
| Liver Sound Speed | Sound Speed Plane-Wave Ultrasound. | Higher |
| Liver Attenuation | Attenuation Plane-Wave Ultrasound coefficient, measuring the decrease in amplitude of the ultrasound waves as a function of frequency. | Lower |
| Liver Elasticity | Elasticity, median of the values measured within the Q-Box. | Lower |
| Shear Wave Velocity | Velocity of the shear wave, median of the values measured within the Q-Box. | Lower |
| Liver Viscosity | Viscosity, median of the values measured within the Q-Box. | Lower |
| Shear Wave Dispersion | Shear wave dispersion median of the values measured within the Q-Box. | Lower |
| **Glycemic Control (CGM)** |  |  |
| Mean Glucose | Mean of all glucose values. | Lower |
| Median Glucose | Median of all glucose values. | Lower |
| GMI | A linear transformation of the mean glucose value, meant to improve the eA1C measure. | Lower |
| Glucose IQR | Interquartile range (IQR), calculated as the distance between the 25th percentile and the 75th percentile of the glucose values. | Lower |
| CV | Coefficient of variation of all glucose values. | Lower |
| MAD | Median Absolute Deviation (MAD). This is a measure of glycemic variability, which is calculated by taking the median of the differences between glucose values and their median. | Lower |
| MODD | Mean difference between glucose values obtained at the same time of day (MODD). This is a measure of glycemic variability that is calculated by taking the mean of the absolute differences between glucose values measured at the same time, a day apart. | Lower |
| MAG | Mean Absolute Glucose (MAG). This is a measure of glycemic variability that’s meant to be less dependent on the mean glucose value and to take into account all variability over time. | Lower |
| SD of ROC | Standard deviation of all the rate of change (ROC) values of an individual. ROC is calculated between every two consecutive glucose measures. | Lower |
| Time-of-Day SD | SD between time points. Standard deviation of the mean glucose value calculated for each time point across all days. | Lower |
| Interday SD | Vertical SD within days. Average value of the SD values for each day separately. | Lower |
| Within-Hour SD | SD within series. Taking hour-long intervals that start at each point of the glucose measures, the SD of each hour-long window is computed, and then the mean of all those SDs is taken. | Lower |
| HBGI | High Blood Glucose Index (HBGI). | Lower |
| LBGI | Low Blood Glucose Index (LBGI). | Lower |
| Time > 140 mg/dL | Percent of glucose measures that were larger than 140 mg/dL. | Lower |
| Time > 180 mg/dL | Percent of glucose measures that were larger than 180 mg/dL. | Lower |
| ADRR | Average daily risk range (ADRR) is a variability measure that was designed to be sensitive to both hyperglycemia and hypoglycemia. The ADRR is composed of the HBGI and LBGI. | Lower |
| COGI | Continuous Glucose Monitoring Index (COGI). COGI uses three measures of the CGM - time in range, time below range and glucose variability (GV) and averages them using a weighted mean, usually with weights of 50%, 35% and 15% accordingly. | Lower |
| GRADE Score | Glycaemic Risk Assessment Diabetes Equation (GRADE). This clinical risk score aims to evaluate the degree of risk presented by a glucose profile. | Lower |
| GRADE Euglycemia | Percentage of the GRADE score that is attributed to euglycemia (glucose values within the normal range, default is 70-140). | Higher |
| **Body Composition** |  |  |
| BMI | Body Mass Index. | Lower |
| Total Fat Mass | Body composition total body fat mass. | Lower |
| Total Body Fat | Body composition total tissue percent fat. | Lower |
| Total Mass | Body composition total tissue mass. | - |
| Waist Circumference | A measure of abdominal fat accumulation. | Lower |
| Hip Circumference | A measure of hip width. | - |
| Waist-to-Hip Ratio | A key indicator of central obesity and metabolic risk. | Lower |
| Android Fat Mass | Body composition android fat mass. | Lower |
| Android Fat % | Body composition android tissue percent fat. | Lower |
| VAT Mass | Total scan visceral adipose tissue (VAT) mass. | Lower |
| VAT Volume | Total scan visceral adipose tissue (VAT) volume. | Lower |
| Trunk Fat Mass | Body composition trunk fat mass. | Lower |
| Trunk Fat % | Body composition trunk tissue percent fat. | Lower |
| SAT Mass | Total scan subcutaneous adipose tissue (SAT) mass. | - |
| SAT Volume | Total scan subcutaneous adipose tissue (SAT) volume. | - |
| Total Lean Mass | Body composition total body lean mass. | Higher |
| Android Lean Mass | Body composition android lean mass. | Higher |
| Trunk Lean Mass | Body composition trunk lean mass. | Higher |

**Table S4:** Counts of Bonferroni-significant associations between oral microbial features and each metabolic phenotype. Positive/negative indicate the sign of the regression coefficient (*β*) from Methods 4.4. This table provides the numeric values for Fig. 1c.

| Phenotype | Strain |  | Gene Family |  | Pathway |  |
| --- | --- | --- | --- | --- | --- | --- |
|  | positive | negative | positive | negative | positive | negative |
| Liver Attenuation | 10 | 7 | 1347 | 35827 | 37 | 32 |
| Liver Sound Speed | 4 | 54 | 2282 | 8275 | 31 | 42 |
| Liver Elasticity | 2 | 6 | 250 | 546 | 1 | 0 |
| Shear Wave Velocity | 4 | 6 | 313 | 440 | 1 | 0 |
| Liver Viscosity | 5 | 0 | 5 | 2451 | 0 | 3 |
| Shear Wave Dispersion | 6 | 0 | 326 | 619 | 3 | 2 |
| Mean Glucose | 19 | 9 | 82680 | 2999 | 109 | 70 |
| Median Glucose | 23 | 8 | 81142 | 2991 | 104 | 64 |
| Glucose IQR | 3 | 23 | 9501 | 2735 | 29 | 29 |
| GMI | 19 | 9 | 82680 | 2999 | 109 | 70 |
| CV % | 4 | 18 | 117 | 5812 | 0 | 15 |
| MAD | 3 | 23 | 12485 | 1864 | 25 | 34 |
| MAG | 11 | 51 | 16314 | 3798 | 13 | 57 |
| MODD | 11 | 12 | 466 | 661 | 0 | 26 |
| SD of ROC | 11 | 72 | 16608 | 4713 | 9 | 63 |
| Time-of-Day SD | 3 | 23 | 21205 | 2124 | 37 | 6 |
| Interday SD | 1 | 18 | 591 | 971 | 0 | 11 |
| Within-Hour SD | 16 | 75 | 32997 | 5295 | 21 | 68 |
| Time > 140 mg/dL | 6 | 8 | 39655 | 2281 | 33 | 55 |
| Time > 180 mg/dL | 0 | 1 | 864 | 44 | 0 | 0 |
| HBGI | 6 | 7 | 42814 | 2514 | 28 | 63 |
| LBGI | 7 | 12 | 1974 | 46929 | 49 | 56 |
| ADRR | 5 | 17 | 1562 | 36234 | 32 | 30 |
| COGI | 7 | 4 | 13601 | 951 | 8 | 17 |
| GRADE Score | 0 | 3 | 2104 | 185 | 0 | 4 |
| GRADE Euglycemia | 2 | 3 | 6623 | 307 | 6 | 2 |
| BMI | 60 | 2 | 5323 | 4204 | 20 | 7 |
| Total Fat Mass | 85 | 2 | 7168 | 5399 | 39 | 9 |
| Total Body Fat % | 115 | 2 | 7774 | 6882 | 38 | 22 |
| Total Mass | 47 | 2 | 5299 | 2664 | 29 | 4 |
| Waist Circumference | 49 | 1 | 7013 | 1625 | 17 | 26 |
| Hip Circumference | 28 | 0 | 2004 | 1064 | 7 | 2 |
| Waist-to-Hip Ratio | 17 | 5 | 8553 | 1004 | 19 | 59 |
| Android Fat Mass | 86 | 2 | 10079 | 3831 | 37 | 28 |
| Android Fat % | 102 | 2 | 9871 | 4105 | 36 | 27 |
| Trunk Fat Mass | 88 | 2 | 9805 | 3850 | 36 | 27 |
| Trunk Fat % | 102 | 2 | 9687 | 4166 | 33 | 28 |
| VAT Mass | 77 | 8 | 19871 | 3874 | 28 | 59 |
| VAT Volume | 77 | 8 | 19871 | 3874 | 28 | 59 |
| SAT Mass | 66 | 2 | 7122 | 3124 | 43 | 5 |
| SAT Volume | 66 | 2 | 7122 | 3124 | 43 | 5 |
| Total Lean Mass | 10 | 4 | 1257 | 436 | 1 | 2 |
| Android Lean Mass | 8 | 7 | 5137 | 1206 | 21 | 4 |
| Trunk Lean Mass | 9 | 7 | 820 | 541 | 1 | 2 |

**Table S5:** Sensitivity analysis excluding participants with pronounced oral problems (*n* = 385). Retention and direction concordance of Bonferroni-significant oral feature– phenotype associations relative to the main analysis (same preprocessing and models; Bonferroni-adjusted *P <* 0.05)

| Oral Feature type | System | Main analysis<br>(no. sig.) | After exclusion<br>(no. still sig.) | Retention<br>(%) | Same direction<br>(%) |
| --- | --- | --- | --- | --- | --- |
| **Strain** | Body | 1,152 | 1,102 | 95.66% | 100.00% |
|  | CGM | 553 | 511 | 92.41% | 100.00% |
|  | Liver | 104 | 89 | 85.58% | 100.00% |
|  | **Total** | **1,809** | **1,702** | **94.09%** | **100.00%** |
| **Gene Family** | Body | 198,749 | 177,553 | 89.34% | 100.00% |
|  | CGM | 592,390 | 492,413 | 83.12% | 96.21% |
|  | Liver | 52,681 | 38,989 | 74.01% | 100.00% |
|  | **Total** | **843,820** | **708,955** | **84.02%** | **97.37%** |
| **Pathway** | Body | 851 | 814 | 95.65% | 100.00% |
|  | CGM | 1,352 | 1,283 | 94.90% | 100.00% |
|  | Liver | 152 | 102 | 67.11% | 100.00% |
|  | **Total** | **2,355** | **2,199** | **93.38%** | **100.00%** |

**Table S6:** Sensitivity analysis adjusting for oral-health scores. Models additionally adjusted for an inflammation score and a periodontal destruction score (alongside age, sex, and smoking status). Retention and direction concordance are reported for Bonferroni-significant oral feature–phenotype associations (Bonferroni-adjusted *P <* 0.05).

| Feature type | System | Main analysis<br>(no. of sig.<br>associations) | After covariate<br>adjustment<br>(no. still sig.) | Retention<br>(%) | Same<br>direction<br>(%) |
| --- | --- | --- | --- | --- | --- |
| **Strain** | Body | 1,152 | 885 | 76.82% | 100.00% |
|  | CGM | 553 | 486 | 87.88% | 100.00% |
|  | Liver | 104 | 63 | 60.58% | 100.00% |
|  | **Total** | **1,809** | **1,434** | **79.27%** | **100.00%** |
| **Gene Family** | Body | 198,749 | 188,056 | 94.62% | 100.00% |
|  | CGM | 592,390 | 589,504 | 99.51% | 100.00% |
|  | Liver | 52,681 | 51,424 | 97.61% | 100.00% |
|  | **Total** | **843,820** | **828,984** | **98.24%** | **100.00%** |
| **Pathway** | Body | 851 | 814 | 95.65% | 100.00% |
|  | CGM | 1,352 | 1,347 | 99.63% | 100.00% |
|  | Liver | 152 | 147 | 96.71% | 100.00% |
|  | **Total** | **2,355** | **2,308** | **98.00%** | **100.00%** |

**Table S7:**
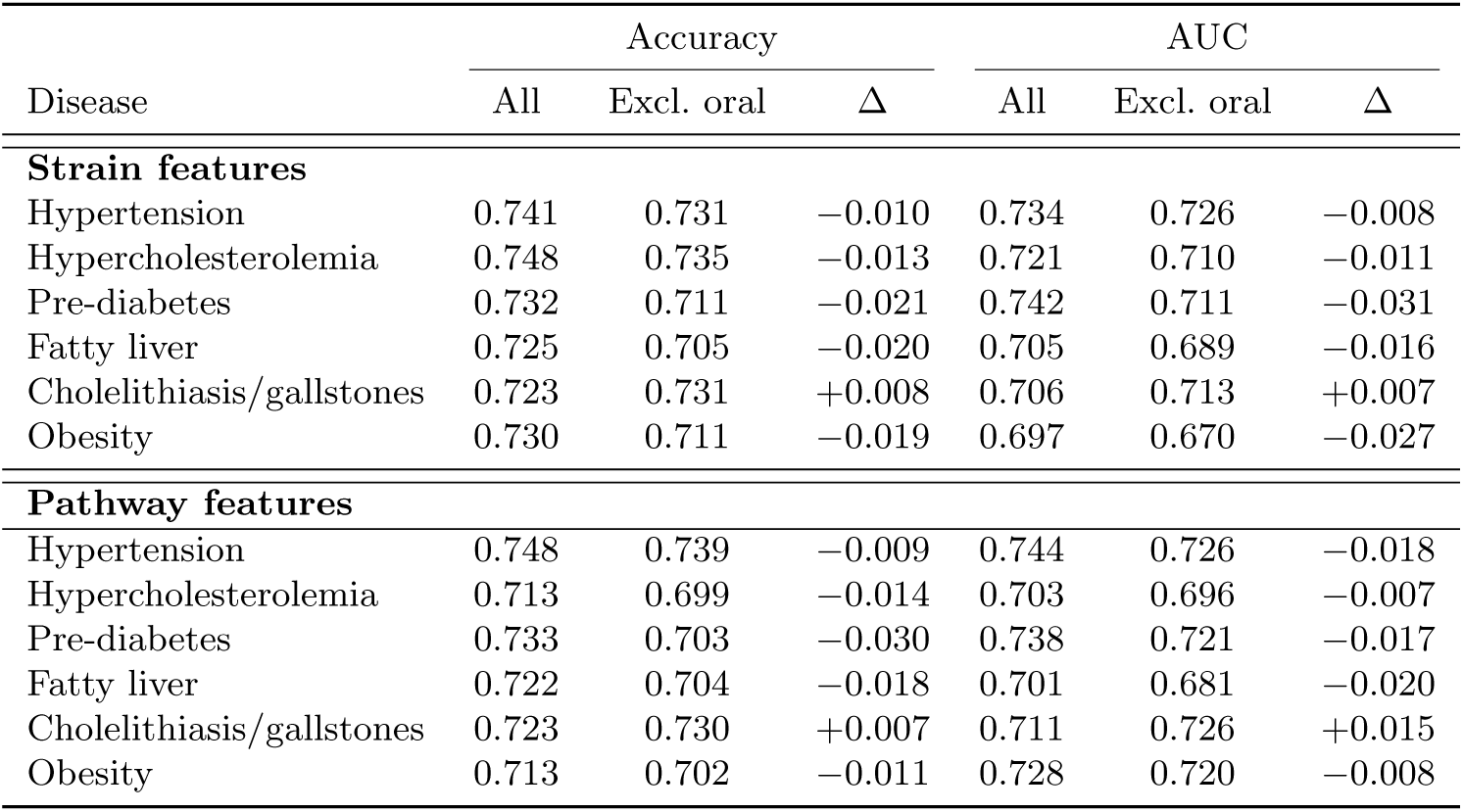
Metabolic disease classification sensitivity analysis after excluding cases and controls with pronounced oral problems. AUC and ACC are reported for models using phenotype-selected oral markers. All: all participants; Excl. oral: excluding participants meeting criteria for pronounced oral problems.

## A Supplementary Results: Community-level contributions to bacterial pathways

To better understand how bacterial communities contribute to functional pathways, we consider two possible modes: (1) a single bacterial taxon encodes the function; (2) multiple taxa collectively contribute to the pathway at the community level. To evaluate these possibilities in pathways associated with metabolic health, we analyzed functional profile data from the HPP dataset. This analysis uses HUMAnN stratified profiles, which decompose each pathway’s abundance into taxon-specific contributions at the genus and species levels. We aggregated contributions to the genus level for clearer visualization. This analysis is descriptive and does not introduce new association results; it is intended to illustrate how the pathway-level functional signals are distributed across taxa in the community.

Among the 299 pathways significantly associated with metabolic health, 9 (3.01%) involve only unclassified bacteria, while the remaining 290 (96.99%) are attributable to known bacterial genera and species. The number of contributing taxa varies across pathways, with an average of 72.20 ± 74.88 species and 26.45 ± 23.82 genera per pathway. Most pathways (275; 91.97%) involve more than one genus, (281; 93.98%) involve more than one species, suggesting that community-level interactions are common.

We further examined the taxonomic composition underlying the 12 most significant microbial pathways (Fig. S1). Favorable-profile pathways exhibited extensive taxonomic contributions, with 55–81 genera and 152–269 species involved per pathway (mean = 66.33 ± 9.33 genera; 211.83 ± 37.85 species). In contrast, adverse-profile pathways involved a much narrower set of taxa, with only 3–7 genera and 4–18 species contributing per pathway (mean = 5.33 ± 2.08 genera; 13.00 ± 7.81 species), suggesting more limited microbial involvement. Pathways showing mixed associations displayed intermediate complexity, with 18–35 genera and 35–72 species per pathway (mean = 25.50 ± 7.94 genera; 57.25 ± 17.35 species).

**Fig. S1:**
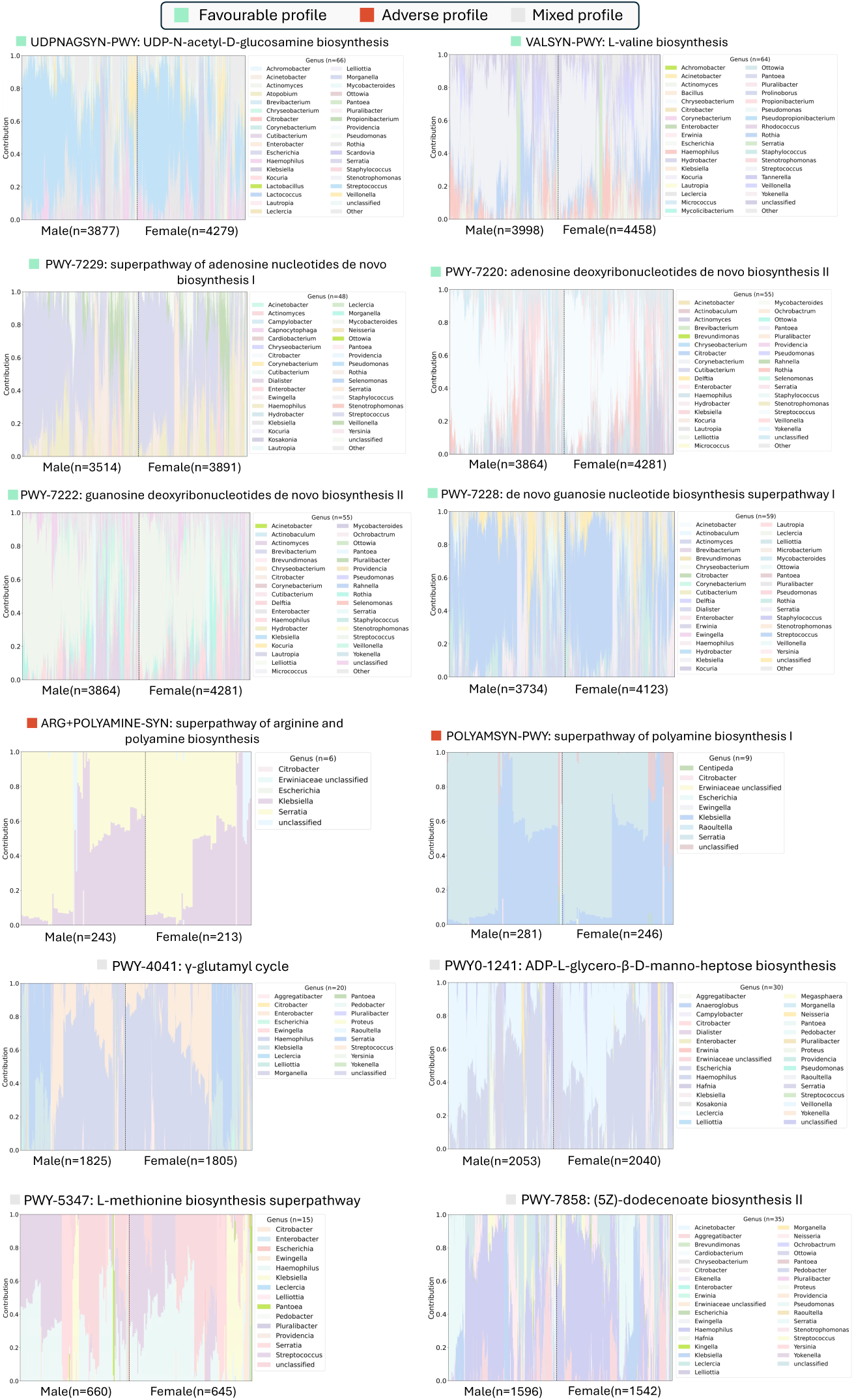
Genus-level contributions to 12 core pathways from Result 2.3.3. Stacked areas show the proportional contribution across male and female participants; only the top contributing genera are shown (remaining grouped as “Other”). The sample size *n* in each panel denotes participants with non-zero abundance and valid genus-level stratification for that pathway, and therefore varies across pathways and sexes. Species-level contributions were aggregated to the genus level for readability. PWY-66-388 is omitted because it is dominated by *Pseudomonas*.

